# Additive in size, synergistic in composition: selection across the eastern oyster larval genome under coastal acidification and sewage effluent

**DOI:** 10.64898/2026.10.08.757713

**Authors:** Jonathan B. Puritz, Johanna A. Harvey, Kathleen E. Lotterhos

## Abstract

Coastal organisms experience natural and anthropogenic stressors simultaneously, yet the interactive effects of multiple stressors are largely unknown. In urbanized estuaries, coastal acidification (CA) is coupled to sewage effluent (SE): nutrient loading from SE stimulates microbial CO_2_ production, and this lowers the pH. The genomic targets of selection of CA and SE have been characterized separately but never together in early life stages of marine organisms. Here, we exposed eastern oyster (*Crassostrea virginica*) larvae to control, CA (∼2800 µatm *p*CO_2_), SE (5% by volume), and the combination (CASE) for 24 h across independent spawns, and used expressed exome capture sequencing to measure allele-frequency change before and after exposure, removing loci that also shifted in controls. CASE generally produced the lowest survival, although mortality was driven primarily by effluent and did not exceed the additive expectation. The size of allele-frequency change under CASE stayed close to the additive sum of the single stressors, but it showed compositional synergy: CASE had more outlier loci than CA and SE combined with 1,941 loci (747 genes) detected only under the combined treatment. CASE outliers were enriched for proteostasis, ciliary function, mitochondrial calcium handling, and a steroid-metabolizing (CYP17) module consistent with effluent endocrine disruptors. Selection acted repeatedly on standing variation across spawns and genetic backgrounds, with the largest allele-frequency shifts at rare, spawn-private loci. Enriched biological functions recurred across spawns more often than expected, while most outlier loci were specific to one or two spawns, consistent with genotypic redundancy. Our results show that combined stressors can recruit a compositionally synergistic genomic response, at loci and functions not detected under either stressor alone, even when survival and the size of allele-frequency change are additive.

**Author summary:** Coastal organisms rarely experience one environmental stress at a time. In urbanized estuaries, coastal acidification occurs alongside sewage effluent, but we know little about how these stressors act together or which genetic variants allow organisms to survive them. We exposed eastern oyster larvae from four independent spawns to coastal acidification, treated sewage effluent, or both for 24 hours. We measured survival in all four spawns and tracked changes in the frequencies of genetic variants in three.

Sewage effluent caused most of the mortality and combining it with coastal acidification increased mortality, but not beyond the additive expectation. The genomic response followed a different pattern. The combined treatment affected more genetic variants than either stressor alone and recruited many variants, genes, and biological functions that we did not detect under either single stressor. Across spawns, selection repeatedly affected the same biological functions, usually through different genetic variants. This result suggests that oyster populations can reach similar biological responses through different genetic routes. The combined treatment selected targets that neither stressor selected alone, so experiments that examine one stressor or one genetic background will miss part of the genomic response that marine organisms face in acidifying, urbanized estuaries.

## Introduction

Coastal marine organisms develop in a multistressor environment, and predicting the combined effect of co-occurring stressors is a major challenge in global-change biology. When two stressors act together their joint effect can be additive, synergistic, or antagonistic, and a synthesis of marine multistressor experiments recovered all three outcomes with no general rule for which to expect [1]. Non-additive interactions are common enough, and damaging enough, to be called ecological surprises, and adding a third stressor changes the interaction in most cases [1–3]. Most experiments still manipulate one stressor at a time and stop at the phenotype, so the mechanisms that produce non-additive outcomes, and whether they leave a signature in the genome, remain largely untested [4,5].

Two stressors that co-occur in urbanized estuaries are coastal acidification and sewage effluent. Nutrient loading from effluent fuels algal blooms, and microbial respiration of that organic matter draws down oxygen while releasing CO_2_ and lowering pH, so hypoxia and acidification are coupled through the same process [6–8]. Sewage treatment outfalls act as point sources of this coupling, and the effluent itself is often acidified and enriched in *p*CO_2_[6]. Effluent also leaves a population-genetic signature: outfalls limit pelagic larval dispersal and reduce gene flow in marine invertebrates whose adults barely move, which places the effect on the larval stage [9,10]. For early life stages of bivalves, hypoxia and acidification combine to depress growth, survival, and metamorphosis in additive to synergistic ways [11,12]. Even so, coastal acidification and sewage effluent have not been studied together as agents of selection.

The eastern oyster (*Crassostrea virginica*) is a foundation species and the basis of a fishery worth close to a billion dollars annually in the United States, and its planktonic larvae are a major survival bottleneck. Larvae are acutely sensitive to acidification during formation of the first shell, when calcification draws on limited maternal reserves and the calcifying surface is exposed to ambient seawater chemistry [13–16]. Larval survival follows a Type III curve, with most individuals dying before settlement, a pattern driven in part by a heavy genetic load of deleterious recessive alleles that environmental stress can expose [17,18]. High fecundity together with high, non-random mortality makes the larval stage a strong and tractable target for selection. Because high genetic load, and the family structure among the larvae in any one cohort, shifts allele frequencies independently of any imposed stressor, reading the genomic response to a stressor depends on separating stressor-driven selection from this background [19].

That selection can be inferred directly from the genome using select-and-resequence experiments. Previous work that has tracked allele-frequency change in pooled larvae across a single generation of stress show that marine invertebrates can adapt rapidly to acidification by reshuffling standing genetic variation, often through polygenic shifts and the recruitment of rare or balanced variants [20–24]. This suggests that the raw material for rapid adaptation is present in wild populations. Where many loci can satisfy the same selective demand, such architectures are redundant, and the responding loci need not be the same ones from replicate to replicate even when the functional response repeats.

What single spawn studies cannot resolve is whether the response is repeatable. Each uses a single pool of larvae from one set of parents, so a consistent adaptive signal cannot be separated from spawn-specific drift or from selection that depends on the particular genetic background [21,22,25]. The difficulty of distinguishing real, replicated allele-frequency change from noise in a single larval cohort is precisely what drove the exchange over temporally balanced selection in Pacific oysters [26–28]. Sweepstakes reproductive success, in which a few parents dominate each cohort, compounds this problem within a spawn but also makes spawns from different parents genuinely independent samples of standing variation [27].

Replicating selection across independent spawns, each from a different set of wild parents, supplies the missing control: loci that respond consistently across genetic backgrounds reflect parallel selection on shared standing variation, whereas background-specific responses point to contingent or polygenic architecture [29].

Here, we address these gaps in a multiple stressor and multiple spawn experiment framework. We exposed eastern oyster larvae to a factorial design crossing coastal acidification and sewage effluent (control, CA, SE, and the combination CASE) for 24 h across independent spawns and used expressed exome capture sequencing [30] of pooled larvae to estimate allele-frequency change before and after exposure against the chromosome-level eastern oyster genome [31,32]. We asked whether CASE produced non-additive effects on larval survival, and whether the genomic response departed from additivity in the size of allele-frequency change, in the identity of the responding loci, or both. We then asked which loci and biological functions the combined treatment recruited that neither single stressor did, how repeatably selection acted across independent genetic backgrounds, and whether recurrent and spawn-specific responses drew on different components of standing variation, partitioned into locus tiers representing different modes of adaptation.

## Results

### Treatment conditions

Treatment conditions were achieved as intended (Table S2). Across the post-exposure bottles, pCO_2_ averaged 542 ± 69 µatm in the controls and 2,815 ± 1,282 µatm under CA, bracketing the nominal 400 and 2,800 ppm set points, with aragonite saturation falling from 1.76 ± 0.33 to 0.56 ± 0.47. Effluent alone raised pCO_2_ only modestly over the control (691 ± 236 µatm; Ω_Ar_ 1.85 ± 0.31), so SE was not itself an acidification treatment, while CASE tracked CA (2,487 ± 623 µatm; Ω_Ar_ 0.61 ± 0.21). Salinity (28.7–29.4) and temperature (19.1–19.3 °C) were comparable across treatments.

### Larval survival

To test whether combined stress killed more larvae than expected from each stressor alone, we measured survival at 24 h in all four spawns (Fig 1C). Survival declined in the order CON > CA > SE > CASE, and this order held under each of four alternative estimates of starting larval density (hereafter density denominators; defined in Methods and in the Fig S2 legend). CASE was the lowest-surviving treatment in every spawn except SP2. A one-way ANOVA on spawn-centered composite survival (12 jars per treatment across four spawns) was significant (*F =* 7.39, *p =* 4.1 × 10⁻⁴). Tukey HSD contrasts localized the effect to SE. CASE survival was 15.3 percentage points below CON (*p =* 0.003) and SE was 14.7 points below CON (*p =* 0.005), while CA sat 2.6 points below CON and did not differ from it (*p =* 0.92). CASE fell 12.7 points below CA (*p =* 0.019) but was statistically indistinguishable from SE (0.6 points, *p =* 0.999).

**Fig 1.**
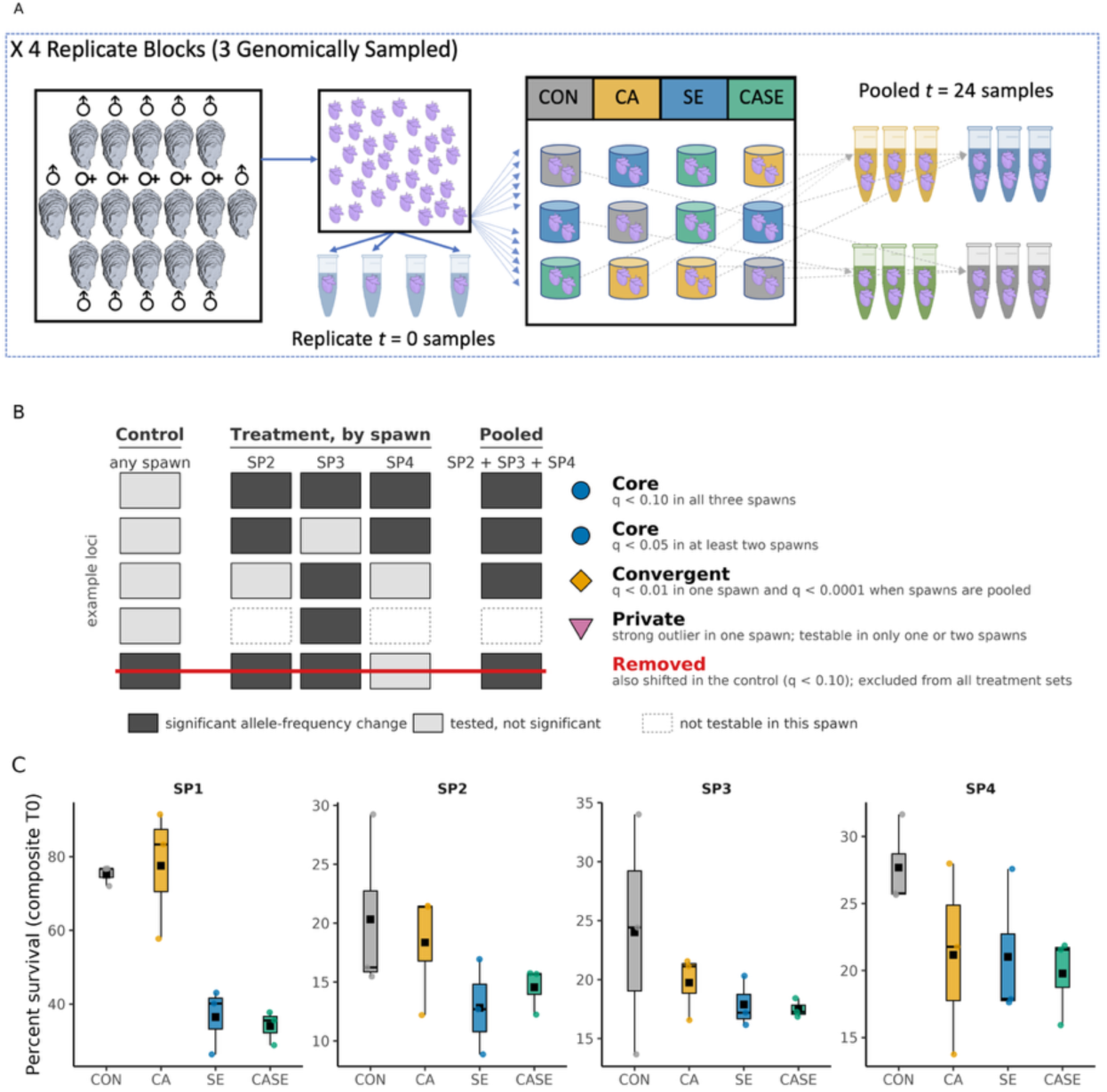
Experimental design, outlier classification, and larval survival. (A) Experimental design schematic. (B) Schematic of how loci were classified (illustrative, not data). Columns show whether a locus changed allele frequency significantly in the control (any spawn), in each sequenced spawn (SP2, SP3, SP4), and in the pooled across-spawn CMH test; dark cells are significant, light cells tested but not significant, dashed cells not testable in that spawn. Loci that also shifted in the control (bottom row) were removed from every treatment set. Remaining outliers were assigned to Core (significant in more than one spawn; circle), Convergent (significant in one spawn and in the pooled test; diamond), or Private (strong outlier in a single spawn; down-triangle); thresholds are given in Methods and the same symbols are used in Figs 4 and 5. (C) Percent survival at 24 h by treatment (CON, CA, SE, CASE) for each phenotyped spawn (SP1, SP2, SP3, SP4); box plots show median and interquartile range with per-jar points and the treatment mean (square), computed on the composite T0 denominator (see Methods; sensitivity to the denominator choice in Fig S2). Survival order CON > CA > SE > CASE, robust across denominators; CASE ≈ SE and CA ≈ CON (Tukey HSD). A factorial CA × SE betabinomial GLMM on composite survival (random spawn effect, all four spawns) found no significant interaction (CA×SE β = 0.04, *p =* 0.88; results/Fig1_survival_GLMM.csv), so combined mortality does not exceed the additive expectation.

The CASE treatment showed no excess mortality beyond the additive expectation. In the high-survival spawn SP1, survival was 75.2% (CON), 77.5% (CA), and 36.5% (SE). Observed CASE survival was 34.1%, just below the additive expectation of roughly 38.8%, a small and non-significant deficit. In the low-survival sequenced spawns SP2 to SP4 (baseline CON around 20% to 28%), observed CASE survival met or slightly exceeded the additive expectation. In SP4, for example, CASE survival was 19.8% observed against 14.5% expected, so the combination did not exceed additive mortality. Larvae under SE and CASE were also more often deformed or dead and, among normal larvae, smaller (Fig S1).

A CA × SE beta-binomial GLMM on composite survival with a random spawn effect, fit across all four spawns, found no significant interaction (CA × SE β = 0.04, *z =* 0.15, *p =* 0.88) while the SE main effect was strong (β = −0.78, *p =* 1.8 × 10⁻⁴). Combined mortality did not exceed the sum of the single-stressor effects, so at the level of survival the response was additive. Per-jar survival under each of the four initial density denominators is shown in Fig S2.

### Sequencing and variant detection

To measure allele-frequency change, we sequenced 57 pooled libraries from paired initial and post-exposure samples across the three 2017 spawns (SP2 to SP4); SP1 was phenotyped only for two reasons. First, the broodstock was only from one population, and second, the genomic sampling differed (tubes and filters) from the other three spawns. Each initial pool contained roughly 40,000 larvae and each post-exposure pool roughly 10,000-20,000 larvae with some control samples having closer to 30,000 larvae. Pools averaged 15.85 ± 5.87 million read pairs of which 99.14% ± 0.18% were retained after trimming and 96.36% ± 3.74% mapped to the *C. virginica* genome (14.04% ± 5.82% duplicates, 90.58% ± 3.39% properly paired). For retained loci, each pool had an average coverage of 59.8 ± 1.42 reads per locus. Within each spawn and treatment, we tested allele-frequency change with the adapted Cochran-Mantel-Haenszel (CMH) test in ACER, using assumed effective population sizes (250 for SP2, 1,000 for SP3 and SP4), the pool sizes above, and the coverage thresholds given in Methods. A locus was called a treatment outlier when its Benjamini-Hochberg false-discovery-rate q-value (hereafter q) in CA, SE, or CASE fell below alpha while its control q exceeded 0.10.

### Population structure across spawns

Two features of oyster larvae shape how allele-frequency change should be interpreted across multiple spawns. Oyster larvae carry high standing variation and a heavy recessive genetic load, so allele frequencies shift during development with no stressor at all, and this load is thought to drive much of early larval mortality in highly fecund bivalves [17,18,33]. Each spawn also draws a different sample of parental variation, and parentage shapes both allele-frequency change and expression in oyster larvae under acidification [28,34]. Both features are in our data. An ordination of 10,000 random loci was dominated by spawn rather than treatment (Fig S3). Additionally, spawns differed in the loci called and in initial allele frequencies, so each is an independent genetic background. We therefore removed any locus that also shifted in the matched control, and we base our inference on outlier signal that replicates across spawns: a locus, gene, or function that recurs across independent spawns is more likely to mark stressor-driven viability selection than the family structure of one cross.

### Reproducibility of outliers across spawns

To ask how repeatable selection was across genetic backgrounds, we compared cross-spawn overlap of treatment outliers with two null models: random loci drawn from each spawn’s own captured set, and a control outlier set built with the same outlier tier rules (Methods). The control shares every technical feature of a spawn, including the probe set, coverage, and the handling response common to all bottles, so it represents a conservative baseline. Loci significant in all three spawns exceeded random expectation for CA (15 loci, 2.1-fold, permutation *p* = 0.006) and CASE (46 loci, 1.3-fold, *p* = 0.05), where the control set fell below it (8 loci, 0.8-fold).

Relative to the control, three-spawn outliers were enriched 2.6-fold under CA (*p* = 0.02) and 1.9- to 2.5-fold under all three treatments when neither set was control-filtered (*p* = 0.01 to 0.03), and a control set filtered reciprocally against treatment outliers retained no three-spawn loci (Fig 2E). Loci significant in at least two spawns exceeded random expectation in every set, including the control (1.14-fold). SE and CASE exceeded the control modestly (1.12- to 1.15-fold), significantly against the full control set (*p* ≤ 0.004) but not against the reciprocally filtered one, which retained only 71 two-spawn loci. Most two-spawn overlap is therefore expected from per-spawn set sizes alone, and the treatment-specific repeatable signal is concentrated in the small set of loci that responded in every spawn.

**Fig 2.**
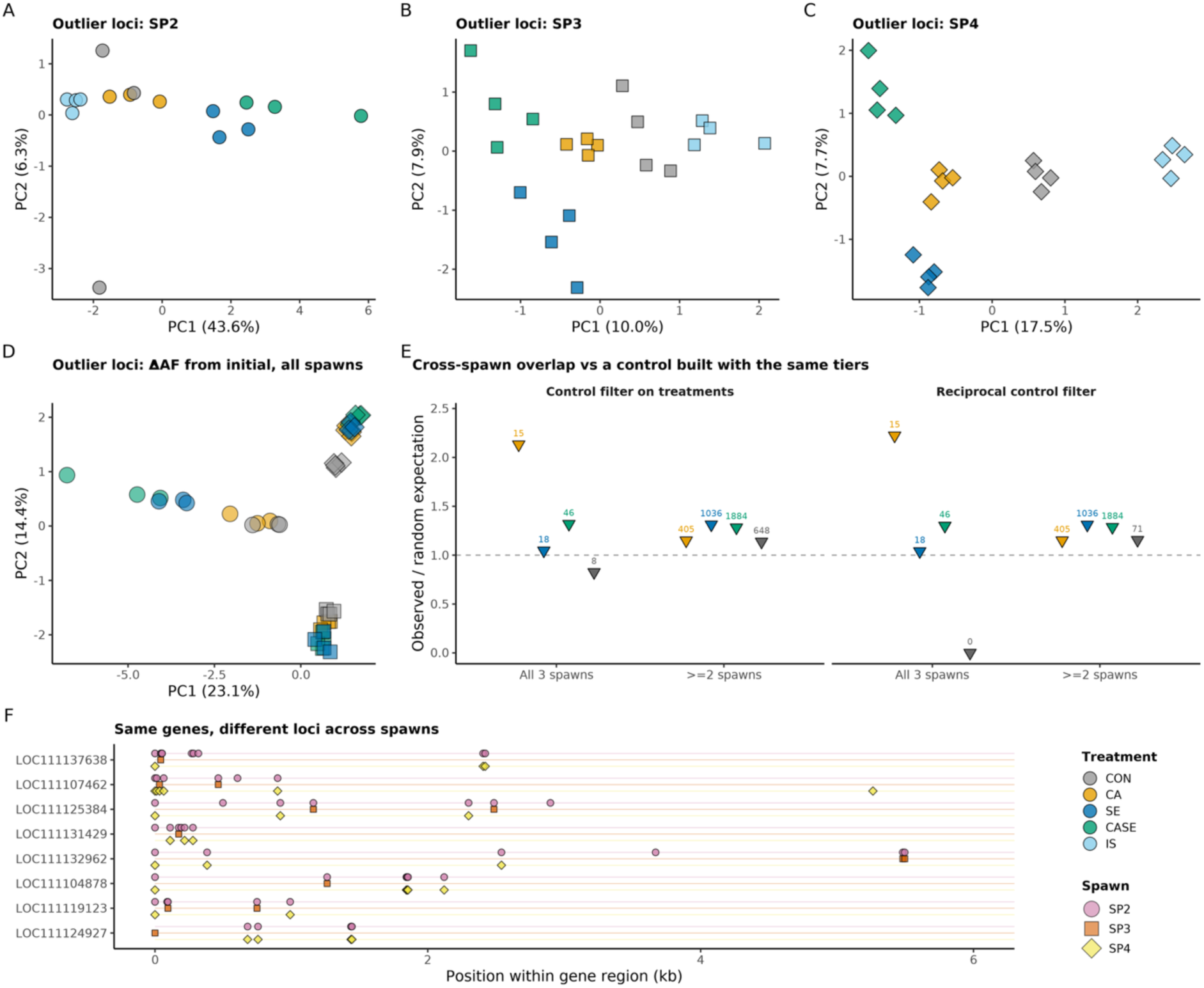
Outlier loci and cross-spawn reproducibility. (A–C) Per-spawn ordinations of outlier loci (SP2, SP3, SP4). (D) Ordination of outlier allele-frequency change from the initial sample across all spawns. (E) Cross-spawn overlap of outlier loci relative to random expectation (dashed line) for CA, SE, CASE, and a control outlier set built with the same tier rules, for loci significant in all three spawns and in at least two; shown with the control filter applied to treatments and with the control filtered reciprocally against treatment outliers. (F) Within-gene locus turnover: recurring genes and the loci that carried them in each spawn; 74 to 89% of recurring genes did so through different loci. Abbreviations: IS, initial (pre-exposure) sample; SP, spawn; CON, control; CA, coastal acidification; SE, sewage effluent; CASE, combined treatment. SP1 was phenotyped only and does not appear in genomic panels.

Genes recurred in all three spawns more often than random genes drawn from each spawn’s captured set (1.02- to 1.07-fold; *p* ≤ 0.03), and 74% to 89% of recurring genes did so through different loci (Fig 2F). Enriched GO terms recurred across spawns more often than terms from random gene sets (terms enriched in all three spawns: 1.37-fold under CA and 1.32-fold under SE, *p* = 0.01; terms enriched in at least two spawns: 1.14- to 1.24-fold for all treatments, *p* ≤ 0.03). The control showed pathway recurrence of similar strength (1.32-fold for terms enriched in all three spawns), and GO-term overlap between different treatments across spawns matched overlap within a treatment, so the functions that recurred were shared across treatments.

### Additivity of allele-frequency change

We next asked whether the size of allele-frequency change under CASE matched the sum of the two single-stressor changes. For each locus we compared the control-anchored CASE change (ΔAF_CASE_ − ΔAF_CON_) with the additive expectation ((ΔAF_CA_ − ΔAF_CON_) + (ΔAF_SE_ − ΔAF_CON_)), anchoring both axes on the control following pool-seq multiple-stressor precedent [35,36]. A slope of 1 means the combined shift equals the sum of the single-stressor shifts. Across all CASE-significant loci the orthogonal slope was 0.99, so in aggregate the combined shift was additive (Fig 3). Departures were confined to subsets. Loci significant only under CASE had a slope of 1.10, with 70% showing changes larger than the additive expectation; this is a modest tendency toward larger-than-additive change, and it sits in the subset most exposed to ascertainment bias. Wherever CA contributed to the significance call (CASE & CA, and CASE & SE & CA) the slope was 0.55, so changes were smaller than the additive expectation. The CA & SE (no-CASE) negative-control panel was flat (slope 0.09).

**Fig 3.**
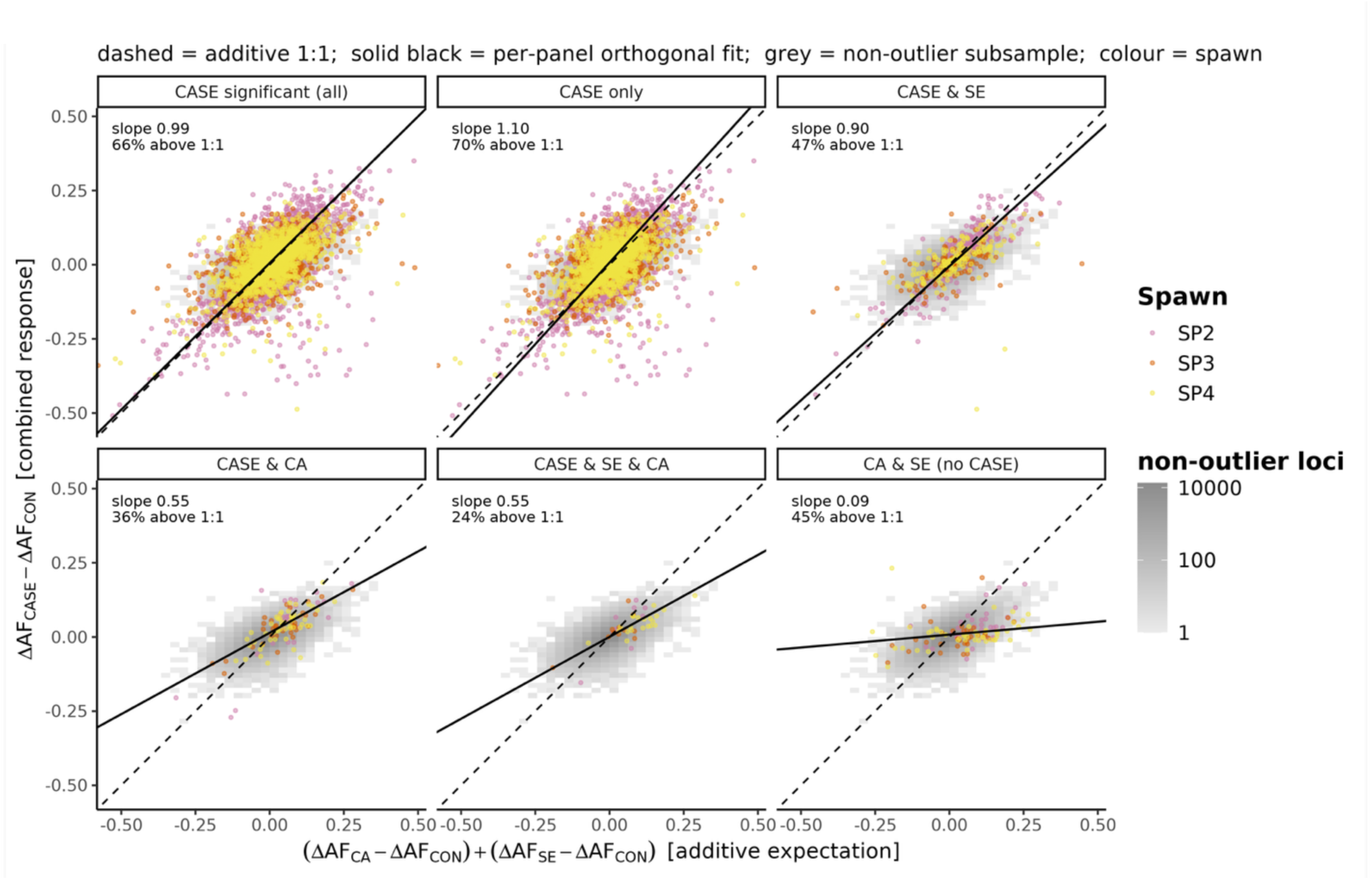
Additivity of allele-frequency change. Control-anchored scatter of the CASE allele-frequency change (y = ΔAF_CASE_ − ΔAF_CON_) against the additive expectation from the single stressors (x = (ΔAF_CA_ − ΔAF_CON_) + (ΔAF_SE_ − ΔAF_CON_)), coverage floor 20. The six panels show different outlier loci subsets: CASE significant (all) is every locus that was significant for the CASE treatment, regardless of significance in other treatments; CASE only, are loci significant in CASE but no other treatment; CASE & SE are loci significant only in the CASE and SE treatment; CASE & CA are loci significant only in the CASE and CA treatment; CASE & SE & CA are loci significant in all three, and CA &SE (no CASE) are loci that are significant both for SE and CA but not for CASE. Grey 2D density is a 10,000-locus subsample of non-outlier loci (the genome-wide additive expectation); colored points are the panel’s loci per spawn; the dashed line is the additive 1:1 expectation and the solid black line the per-panel orthogonal (Deming) fit, with the orthogonal slope and the percentage of loci above the 1:1 line annotated per panel (one point per locus, averaged across spawns, in the fit). Slopes near 1 indicate additivity. Changes smaller than the additive expectation where CA contributes are the robust pattern; the modest tendency toward changes larger than the additive expectation among CASE-only loci should be read cautiously, because outlier ascertainment pushes points upward. The CA & SE (no CASE) panel is a negative control.

### Treatment-exclusive loci

We then asked whether CASE selected the same loci as the single stressors or recruited its own. We call a locus treatment-exclusive when it is a significant outlier in one treatment and in neither of the other two (analogous to a private allele, but for outlier status). If the CASE response were built only from the CA and SE responses, CASE-significant loci should fall inside the union of the two single-stressor sets. Across all spawns, exclusive locus counts were 375 (CA), 761 (SE), and 1,941 (CASE), with only small pairwise overlaps (64 CA and SE, 70 CA and CASE, 232 SE and CASE) and 29 shared by all three (Fig 4C). The CASE-exclusive set exceeded the CA- and SE-exclusive sets combined (1,136), and the same held at the gene level (747 CASE-exclusive genes against 143 for CA and 291 for SE). We refer to this pattern as compositional synergy: the combined treatment recruited more exclusive loci and functions than the two single-stressor responses together, even though the size of allele-frequency change stayed additive. Because significance thresholds and detection power determine which loci are called exclusive, the term describes the composition of detected targets rather than a demonstrated absence of response under the single stressors. Tier-classified outliers were distributed across the genome in all three stressors (Fig 4A for SP4; Fig S7, S8 for SP2 and SP3).

**Fig 4.**
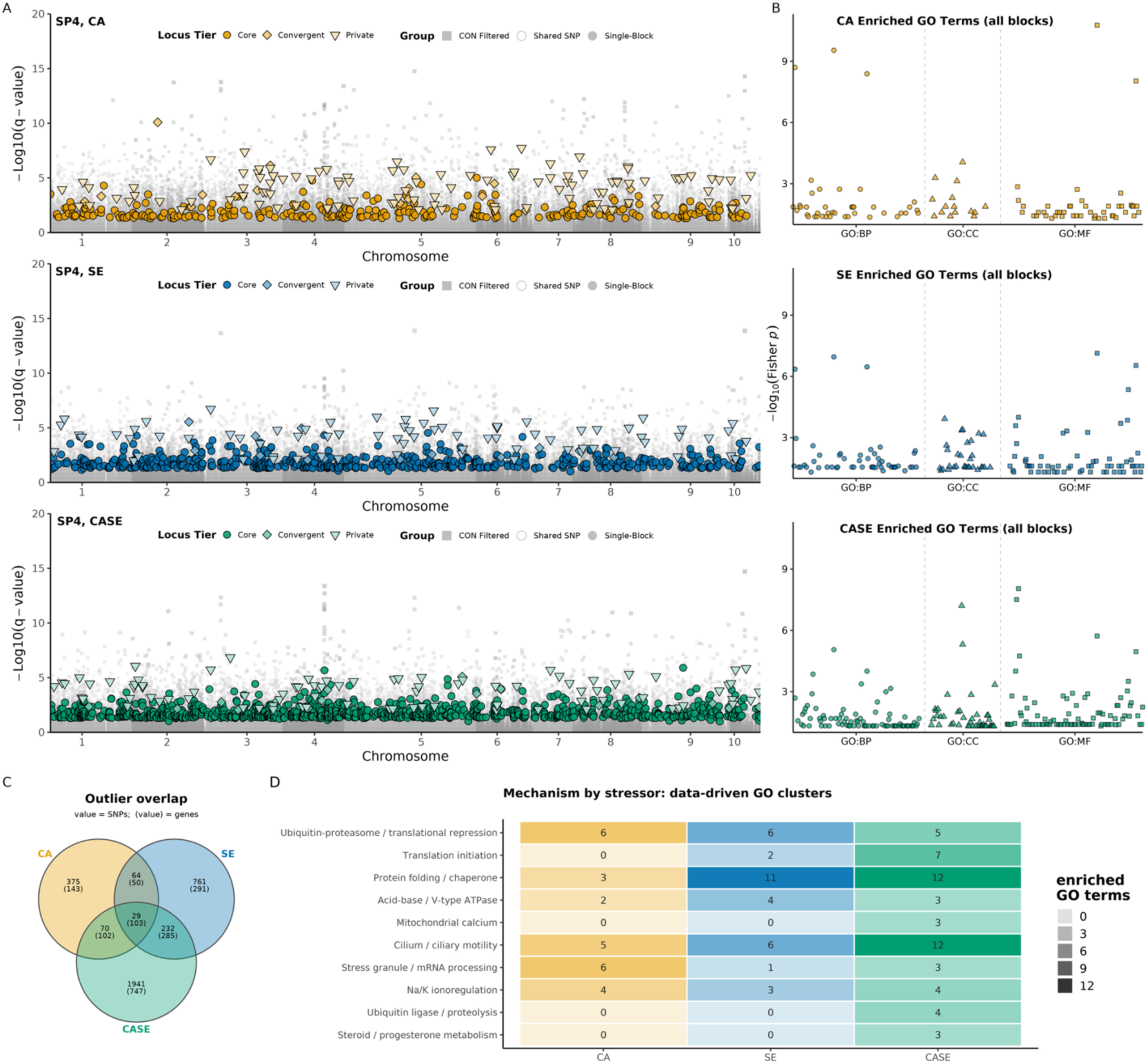
Treatment-exclusive loci and functional enrichment. (A) Manhattan stack for SP4, one row per stressor (CA, SE, CASE), plotting -log10(q-value) against chromosome (1-10); background loci shaded by signal strength and grouped as shared (significant in more than one spawn), single-spawn (significant in this spawn only), or control-filtered (removed because they also shifted in the control); tier-classified outliers (Core, Convergent, Private; defined in Fig 1B and Results, Locus tiers) overlaid as filled symbols (Core circle, Convergent diamond, Private down-triangle) in the per-stressor color; note that the y-axis is capped at 20 (two control-filtered points on chromosome 5 above the cap are noted, not shown). (B) GO enrichment by stressor (topGO): one point per enriched GO term, ordered by GO category (BP, biological process; CC, cellular component; MF, molecular function) with dashed separators, y = -log10(Fisher p), one row per stressor. (C) Outlier overlap among CA, SE, and CASE as a Venn in which each region carries the locus count (top number) and gene count (in parentheses); the CASE-exclusive set is the largest and exceeds the additive expectation from CA and SE. (D) Stress-relevant GO clusters (rows) by stressor (columns), tile shading scaled to the number of enriched GO terms; several clusters are CASE-specific. Clusters were defined by gene-overlap community detection (Methods).

### Gene ontology enrichment

To identify the biological functions under selection, we tested outlier genes for Gene Ontology (GO) enrichment within each treatment. All three treatments were enriched for genes involved in proteostasis. CASE showed strong enrichment for ubiquitin ligase activity, proteasome-mediated catabolism, translational repression, and ribosomal function (Fig 4D; Table 1). CASE contained 212 uniquely enriched GO terms, compared with 102 under CA and 144 under SE, and 153 in the combined analysis. Normalized per 100 candidate genes these were 36.2 (CA), 25.8 (SE), and 22.5 (CASE), so the absolute rise in CASE terms did not track candidate-gene number. The strongest CASE enrichments were ubiquitin protein ligase activity (*p =* 1.1 × 10⁻⁵, 47 genes), proteasome-mediated ubiquitin-dependent catabolism (*p =* 1.0 × 10⁻⁴), protein polyubiquitination, translation repressor activity (*p =* 1.9 × 10⁻⁶), and negative regulation of translation (*p =* 8.8 × 10⁻⁶). Terms detected only under CASE were led by structural constituent of ribosome (*p =* 9.0 × 10⁻⁹, 26 genes), mRNA binding (*p =* 3.1 × 10⁻⁸), cytosolic large ribosomal subunit (*p =* 6.2 × 10⁻⁸), and translation initiation factor activity (*p =* 1.8 × 10⁻⁵).

**Table 1.** Data-driven GO clusters. Enriched GO terms grouped by the overlap of their candidate genes (Jaccard index at least 0.10), using greedy-modularity community detection on the term-similarity network; clusters with at least three terms are reported. For each cluster the table gives the number of enriched terms per stressor (CA, SE, CASE), the stress-relevant flag, the minimum Fisher p across member terms, and representative terms. Member genes for every cluster are in Table S7. Stress-relevant clusters are the ones shown in Fig 4D. The clustering is data-driven; the stress designation and cluster names are an interpretive overlay.

| Cluster | CA | SE | CASE | Terms | Min Fisher | CASE-only | Representative terms |
| --- | --- | --- | --- | --- | --- | --- | --- |
| Ubiquitin-proteasome / translational repression | 6 | 6 | 5 | 6 | 1.70e-11 | no | translation repressor activity; negative regulation of translation; protein polyubiquitination |
| Translation initiation | 0 | 2 | 7 | 7 | 3.10e-08 | no | mRNA binding; translation initiation factor activity; translation reinitiation |
| Protein folding / chaperone | 3 | 11 | 12 | 15 | 4.40e-06 | no | unfolded protein binding; chaperonin-containing T-complex; protein folding |
| Acid-base / V-type ATPase | 2 | 4 | 3 | 4 | 2.00e-04 | no | proton-transporting ATPase activity, rotational mechanism; proton transmembrane transport; vacuolar proton-transporting V-type ATPase complex |
| Mitochondrial calcium | 0 | 0 | 3 | 3 | 4.20e-04 | yes | calcium import into the mitochondrion; uniplex complex; mitochondrial calcium ion homeostasis |
| Cilium / ciliary motility | 5 | 6 | 12 | 16 | 5.10e-04 | no | acrosomal vesicle; ciliary basal body; cilium movement involved in cell motility |
| Stress granule / mRNA processing | 6 | 1 | 3 | 8 | 6.80e-04 | no | mRNA splice site selection; cytoplasmic stress granule; poly(A) binding |
| Na/K ionoregulation | 4 | 3 | 4 | 4 | 1.86e-03 | no | cellular sodium ion homeostasis; cellular potassium ion homeostasis; sodium ion export across plasma membrane |
| Ubiquitin ligase / proteolysis | 0 | 0 | 4 | 4 | 1.28e-02 | yes | ubiquitin protein ligase binding; modification-dependent protein catabolic process; regulation of proteolysis |
| Steroid / progesterone metabolism | 0 | 0 | 3 | 3 | 2.21e-02 | yes | steroid 17-alpha-monooxygenase activity; 17-alpha-hydroxyprogesterone aldolase activity; progesterone metabolic process |

We also asked whether the two single stressors selected on the same functions. The CA and SE signatures were functionally distinct. CA enriched for transcript buffering and metabolic tuning (cytoplasmic stress granules, mRNA 3’-UTR binding, RNase MRP, glycolytic and redox enzymes). SE enriched for protein-folding capacity and cytoskeletal and cell-cycle remodeling (chaperonin TCP, nascent polypeptide-associated complex, cortical actin, mitotic cycle). The overlap was asymmetric: SE and CASE shared 47 enriched terms whereas CA and SE shared only 16, so the combined response drew more heavily on the SE program, and a steroid and progesterone cluster (exogenous-steroid handling) was specific to CASE.

Grouping enriched terms into clusters by the candidate genes they share (community detection on gene overlap; Methods) (Fig 4D, Table 1) recovered proteostasis (ubiquitin-proteasome with translational repression, minimum Fisher *p =* 1.7 × 10⁻¹¹) and protein folding and chaperone function (15 terms, weighted toward SE and CASE) as the strongest modules, together with cilium and ciliary motility (16 terms). Three clusters were CASE-specific: mitochondrial calcium import (3 terms), ubiquitin ligase and proteolysis (4 terms), and steroid and progesterone metabolism (3 terms). Translation initiation was also concentrated in CASE (7 of its terms). The combined stressor engaged bioenergetic and protein-homeostasis machinery that neither single stressor did alone.

### CASE-specific gene sets

We looked closely at CASE-specific gene sets because they mark non-additive responses to combined stress that survival data alone would miss. Four enriched terms detected only under the CASE treatment resolved to small, named gene sets. Calcium import into the mitochondrion (GO:0036444, *p =* 4.2 × 10⁻⁴, with the uniplex term GO:1990246) resolved to the mitochondrial uniporter holocomplex and formed a CASE-only cluster together with mitochondrial calcium ion homeostasis (Table 1; member genes for all clusters in Table S7). Calcium-dependent cysteine-type endopeptidase activity (GO:0004198, *p =* 3.5 × 10⁻³, 5 of 26 annotated genes) resolved to five calpain genes (calpain-5, calpain-8, calpain-9, and an androglobin/calpain-7-like gene) and did not join any cluster. Ubiquitin-dependent ERAD (GO:0030433, *p =* 7.5 × 10⁻³, 7 of 48 annotated genes) resolved to the core ERAD machinery (VCP/p97, BiP/GRP78, calnexin, calreticulin, and selenoprotein S) and fell within the protein folding and chaperone cluster shared across treatments. Steroid handling (steroid 17-alpha-monooxygenase GO:0004508, *p =* 0.022; 17-alpha-hydroxyprogesterone aldolase GO:0047442; progesterone metabolic process GO:0042448) resolved to two tandem CYP17A1-like paralogs encoding steroid 17-alpha-hydroxylase/17,20-lyase, and likewise formed a CASE-only cluster.

### Locus tiers

To separate loci that respond the same way in every genetic background from loci whose response depends on the background, we assigned outliers to one of three tiers by how their signal replicated across spawns (schematic in Fig 1B; thresholds in Methods). Core loci were significant on their own in more than one spawn (*q* < 0.10 in all three spawns, or *q* < 0.05 in at least two): the replicated, background-independent class. Convergent loci were significant in at least one spawn and again when all three spawns were combined in a single across-spawn CMH test, but did not meet the Core criteria: weak, consistent shifts that only the pooled test resolves. Private loci were strong outliers in a single spawn among loci testable in only one or two spawns (top 1% of that treatment’s *q*-distribution, with spawn-specific thresholds): the background-dependent class. CASE dominated every tier (Table S3): Core 2,436 loci (326 CA, 822 SE, 1,479 CASE), Convergent 493 (31, 83, 406), and Private 543 (178, 175, 370).

### Allele-frequency architecture by tier

We then asked whether the three tiers drew on different components of standing variation, comparing starting minor-allele frequency and the size of allele-frequency change across tiers (Fig 5). Core loci were common, with higher starting minor-allele frequency and modest, consistent shifts, indicating parallel selection on standing variation. Private loci were rarer and carried larger frequency changes, a background-dependent class with larger shifts. Convergent loci were individually weak and polygenic, emerging only when spawns were pooled. These patterns were consistent across spawns (Fig S16).

**Fig 5.**
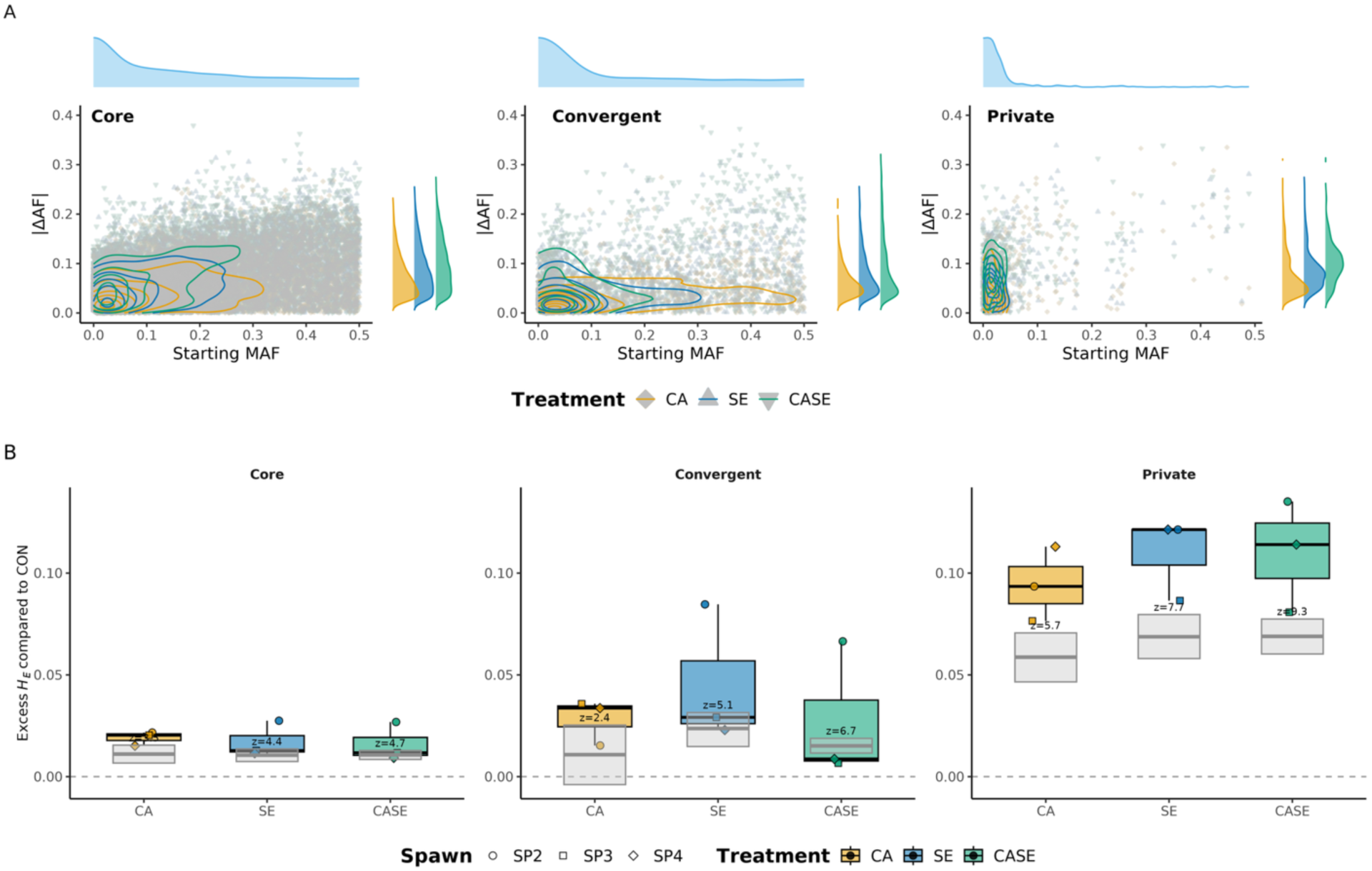
Diversity, architecture, and convergent function of the locus tiers. (Top) For each tier (Core, Convergent, Private; defined in Fig 1B), the two-dimensional kernel density of starting minor allele frequency (MAF0, x) against the magnitude of allele-frequency change (|ΔAF|, y), colored by stressor, with the MAF0 density (top margin) and a per-stressor |ΔAF| ridgeline (right margin); control excluded. Core loci are common with modest shifts, Private loci rarer with larger shifts, Convergent loci a weak polygenic signal. (Bottom) Diversity change at selected loci by tier: for each treatment, the excess change in expected heterozygosity at its outlier loci (the treatment’s post-minus-initial He change minus the control’s change at the same loci) for the Core, Convergent, and Private tiers (points are spawns), with the frequency/shift-matched neutral-null band overlaid (grey) and the z of the observed excess above the null annotated per tier. Positive values indicate diversity raised at selected loci relative to control, and where the excess clears the null band, above the matched-null expectation; magnitudes are not genome-wide estimates because loci are ascertained on allele-frequency change.

### Functional enrichment by tier

GO enrichment also differed among tiers, so the functions under selection tracked how repeatably a locus responded (Fig S15). Core loci were enriched for proteostasis (translation repressor activity *p =* 1.7 × 10⁻¹⁵, negative regulation of translation *p =* 3.8 × 10⁻¹⁴, ubiquitin protein ligase activity *p =* 2.4 × 10⁻¹³, proteasome-mediated catabolism *p =* 3.2 × 10⁻¹²).

Convergent loci were enriched for energy metabolism and signaling (adenylate kinase activity *p =* 1.6 × 10⁻⁴, GTPase activity *p =* 5.1 × 10⁻⁴, motile cilium *p =* 1.7 × 10⁻³, steroid 17-alpha-monooxygenase *p =* 1.8 × 10⁻³). Private loci were enriched for cytoskeletal and chaperone functions (actin filament binding *p =* 2.4 × 10⁻⁵, unfolded protein binding *p =* 1.0 × 10⁻⁴, axonemal dynein complex assembly *p =* 1.4 × 10⁻⁴, cortical actin cytoskeleton *p =* 1.8 × 10⁻⁴).

### Diversity at selected loci

Finally, we asked what selection did to diversity at its targets. A classic sweep drives one allele toward fixation and lowers heterozygosity, whereas a rare allele that rises in frequency without fixing raises it, so the direction of change at outliers separates the two. Selection raised diversity at its own targets. For each treatment, we measured the change in expected heterozygosity (post minus pre) at that treatment’s outliers and subtracted the same change at control loci. The excess was positive in every tier and treatment. To confirm that this exceeded ascertainment and drift, we compared each excess with a frequency- and shift-matched neutral null, and the observed excess cleared the matched-null band in every tier (z from about 2.4 to 9.3, marginal only for Convergent CA; Fig 5, Fig S14). The excess scaled from about 0.016 to 0.019 (Core) to

0.027 to 0.046 (Convergent) to 0.094 to 0.110 (Private), a 5- to 6-fold rise from common Core loci to rare Private loci that held across all three spawns. Because loci were ascertained on allele-frequency change, we base our interpretation on the positive direction and the Core-to-Private gradient. Selection pushed low-frequency standing variants toward intermediate frequency, most strongly among the rare Private loci that carried the largest shifts.

## Discussion

We used a factorial design to ask how the eastern oyster genome responds to coastal acidification, sewage effluent, and their combination across independent wild spawns. Mortality was driven mainly by effluent: acidification alone tracked the controls, and the CASE treatment was statistically indistinguishable from effluent alone. While at the level of survival the stressors acted additively, our results showed evidence of compositional synergy at the genomic level.

The size of allele-frequency change under CASE stayed close to the additive sum of the single-stressor changes, but the combined treatment recruited 1,941 loci, 747 genes, and several biological functions that neither single stressor did, more than both single-stressor-exclusive sets combined. This compositional response is our evidence for compositional synergy.

Selection acted on standing variation, with common alleles shifting modestly and rarer alleles shifting more, and it raised heterozygosity at its targets. Across spawns the same genes and functions were selected through different variants, so functions were more repeatable across genetic backgrounds than individual loci, the pattern expected under genotypic redundancy.

### Multistressor larval mortality was additive

At the level of survival, the combined-stressor response was additive and driven mainly by effluent. CASE was statistically indistinguishable from SE alone, and CA by itself never separated from control. The lack of a detectable acidification effect on survival contrasts with studies in which marine invertebrate larvae raised at moderately elevated *p*CO_2_ for days to weeks showed elevated mortality, shell deformities, delayed metamorphosis, and slower growth [13–15,37–40]. Our exposure was 24 h at high *p*CO_2_, closer to the short, intense acidification pulses estuarine oysters meet in nature [41,42], and eastern oyster larvae can grow at *p*CO_2_ levels well above those used here [43].

The observation that SE induces larval mortality is consistent with population-level studies showing effects of effluent on population connectivity. Puritz and Toonen surveyed 16 populations of a non-harvested species of sea star with a high dispersal potential across the Southern California Bight, and used a multivariate seascape model [44,45] to show that SE was an effective barrier to dispersal, limited geneflow, and that populations closest to effluent sources have lower genetic diversity [9]. Recent work in mussels in the Baltic Sea has found a similar pattern of sewage effluent limiting gene flow among populations [10]. For both sea stars and mussels, adult migration is extremely limited, so sewage effluent must be acting as a barrier to dispersal of pelagic larvae in the water column.

Others have reported synergistic mortality under paired acidification and low-oxygen or contaminant stress in early-stage bivalves [11,12]. We did not detect it, and our design may have lacked the power to do so: baseline mortality in three of four spawns was already high, which compresses treatment differences, and small-volume density counts may lack the precision to resolve a modest interaction. Additive survival outcomes are nonetheless common in pairwise stressor experiments, roughly as common as synergistic or antagonistic ones [1,5,46].

### The size of allele-frequency change under CASE was additive

Ours is among the first factorial designs to test whether the size of allele-frequency change under combined stress is additive, and in aggregate it was. Across CASE outlier loci the combined shift was well predicted by the sum of the CA and SE shifts (slope 0.99). Departures were confined to subsets. Loci significant only under CASE showed changes somewhat larger than the additive expectation, but this is also the subset most exposed to ascertainment bias, since conditioning on CASE significance preferentially retains larger combined shifts. Loci significant under CASE and CA but not SE showed changes smaller than the additive expectation, consistent with SE acting antagonistically at some acidification targets.

Earlier pool-seq studies of larvae under two conditions did not test additivity directly, but they did find hints of non-additive genomic responses. Brennan et al. (2019) found the largest allele-frequency shifts in urchin larvae at loci shared between pH regimes [22], and two multistressor studies inferred combination effects from the size of the shift under the harsher condition [47] or from the recruitment of new targets [35,36]. Measured against a formal additive sum, our result is that the size of change is additive, matching the broad expectation that interaction strength is often modest even when two stressors are individually potent [1,5,46].

### CASE selected loci and functions detected under neither single stressor

Compositional synergy was the non-additive part of the response: CASE selected loci that neither single stressor did. If CASE was simply a composite of the single stressors, CASE-significant loci would have fallen inside the union of the CA and SE sets. Instead, CASE recruited 1,941 loci and 747 genes detected under neither single stressor, more than the CA- and SE-exclusive sets combined. Some of this recruitment is expected even under additivity: for about three-quarters of CASE-exclusive loci, CA and SE shifted the same allele in the same direction, but too little to be detected alone, and the combined exposure pushed them past detection. Compositional synergy therefore describes what the combined treatment makes visible to selection and to us, not an interaction at each locus; a minority of CASE-exclusive loci (∼14%) showed CASE shifts with little single-stressor precedent. This is the type of response Kroeker et al. (2017) anticipated in arguing that coping with acidification amid co-occurring stressors would engage physiology beyond the sum of the parts [46]. Environment-specific outlier sets are a recurring feature of single-stressor marine larval studies; Bitter et al. (2019) found 58% of their low-pH outliers unique to that environment [21], and copepods under warming and acidification recruited new targets under a two-stressor design [35,48]. A factorial design adds the direct test of whether the combination exceeds the single-stressor union, and we convincingly observed this pattern.

This compositional synergy was superimposed on a response common to all three treatments. CA, SE, and CASE shared 20 GO terms: ubiquitin-proteasome degradation, translational repression, acid-base regulation, and ion homeostasis. Acute cellular stress in larvae should involve this type of response, and it maps onto the conserved proteostatic machinery that oysters deploy under environmental stress, including an unfolded protein response, ER-associated degradation, and an expanded set of chaperones and inhibitors of apoptosis. Ubiquitin-pathway variants are recurrent targets of pH-related selection in other marine invertebrates as well [23]. CASE then added terms that were not significant in either single stressor.

Hypoxia-plus-acidification frameworks predict that when two stressors draw on one larval energy pool, aerobic capacity becomes the shared bottleneck [49,50]. Several CASE-exclusive modules fit that prediction and line up into one physiological route: ER stress coupled to mitochondrial calcium loading, calpain activation, protein quality control, and a drop in aerobic capacity. The mechanistic links here are established outside molluscs: ER stress drives mitochondrial calcium overload and a complex I defect [51], mitochondrial calpain is activated during ER stress and under hypoxia-linked oxidative stress [52,53], and proteasome or UPS perturbation is itself a potent trigger of the cellular stress response [54,55]. The genes under those terms include MCU/MICU calcium handling, calpains, and the ERAD machinery p97/VCP, BiP, calnexin, calreticulin, and selenoprotein S. We interpret this as an energy-budget effect: acidification raises the acid-base and calcification cost during the brief early-larval window when reserves are largely maternal [14,15]. Sewage effluent adds contaminant and folding stress, and only when both act together does this bioenergetic pinch point show up in the genome.

The second CASE-only module, steroid metabolism, indicates the impacts of effluent chemistry. Steroid 17-alpha-monooxygenase, 17-alpha-hydroxyprogesterone aldolase, and progesterone metabolism enriched only under CASE and recurred across spawns, carried by two tandem CYP17-like cytochrome P450 paralogs. Molluscs do not appear to run a classical vertebrate sex-steroid axis: their genomes lack the cholesterol side-chain cleavage enzyme CYP11A1 and aromatase CYP19A1, their estrogen receptors are constitutively active and do not bind estrogen, and the case for de novo vertebrate-steroid synthesis is weak [56–60]. Molluscs do, however, take up waterborne steroids from the environment with high capacity and esterify and store them, a route consistent with selection on uptake, transformation, or detoxification of effluent-borne steroids rather than on an endogenous hormone axis [61–63]. Sewage effluent is a well-characterized source of estrogens and other endocrine-disrupting compounds that impair bivalve and echinoderm early development at environmentally relevant concentrations [64–66]. That this signal surfaced only under CASE suggests acidification changed how larvae handled effluent-borne steroids, perhaps through the same shared energy budget.

### Functions repeated across spawns while variants did not: genotypic redundancy

Most outlier loci were specific to one or two spawns, and only a small set responded in all three more often than expected, the low locus-level repeatability that high genotypic redundancy predicts. Repeatability beyond chance extended to higher biological levels: genes recurred in all three spawns slightly more often than random genes, mostly through different loci, and enriched GO terms recurred across spawns 1.2- to 1.4-fold more often than terms from random gene sets. This is genotypic redundancy: when many variants can move a shared function, no single one is required, and the identity of the responders varies from replicate to replicate even as the functional response repeats [67–70]. The control showed pathway recurrence of similar strength, and treatments resembled each other across spawns as closely as themselves, so the repeatable functional core reflects a response to stress and development shared across treatments; stressor specificity lies in the treatment-exclusive modules described above.

Genetic redundancy was different across the three different tiers of outlier loci. Core loci, those with signal in two or more spawns, were more repeatable: common variants with moderate, consistent shifts, enriched for the same functions of proteostasis and translational repression. Private loci, significant in one spawn only, were not repeatable: rarer variants with larger shifts confined to a single spawn, enriched for background-specific processes such as actin filament binding, unfolded protein binding, and axonemal dynein. Convergent loci fell between, significant in at least one spawn and again when all spawns were pooled, enriched for energy and signaling functions including adenylate kinase and GTPase activity, motile cilia, and the steroid 17-alpha-monooxygenase signal. Repeatability declined as variants became rarer and shifts became larger, and the functions engaged moved from a shared core toward spawn-specific processes. A single-pool experiment can name selected loci but cannot tell a portable target from one background’s route to a shared function, which is why detecting parallel polygenic adaptation requires replicated backgrounds [71,72].

Effluent was collected the day before each spawn and likely differed in composition, whereas CA chemistry was held constant. CA, the only treatment applied identically in every spawn, had the strongest three-spawn signal, and because SE and CASE were not perfect environmental replicates across spawns, their cross-spawn recurrence is a conservative measure. The CASE-exclusive modules (mitochondrial calcium handling, calcium-activated proteolysis, ER-associated degradation, steroid processing) carry the stressor-specific part of the response, layered on the functional core shared across treatments.

Genetic drift would be the alternative hypothesis to explain our results but does not fit the data well. CASE, on average, killed the most larvae, leaving the smallest survivor pools and the widest opening for sampling noise. However, several lines of evidence do not support the drift hypothesis. For one, the CMH tests incorporated an assumed effective population size and a conservative post-exposure pool size (Methods) [73,74]. Secondly, loci responding in all three spawns exceeded both random expectation and a control set built with identical rules, and, lastly, functional themes recurred across independent spawns. Drift would scatter signal across functions as well as loci, so functional recurrence paired with locus turnover best fits non-parallel selection on a redundant architecture.

The genomic response was also decoupled from our fitness proxy, survival. CA produced reproducible allele-frequency shifts across spawns with no detectable drop in survival relative to controls, and CASE produced the largest genomic response with survival no different from SE alone. Selection therefore reshaped allele frequencies below the resolution of aggregate mortality.Decoupling of survival from the genomic signature has been reported before in other larval selection experiments. Bitter et al. (2019) reared *Mytilus galloprovincialis* larvae at ambient and low pH, recovered near-identical mortality in both (about 97.7%), and still identified distinct sets of selected loci in each environment [21]. Brennan et al. (2019) reported the same in purple urchin larvae: mortality at pH 8.0 and pH 7.5 differed by less than a point (50.8 versus 51.5%, not significant) while the genetic targets of selection diverged [22]. This apparent decoupling between the genomic response to selection and population-level survival is consistent with genotypic redundancy: many genotypes reach the same fitness, so which larvae survive can change without changing how many do [70].

### Variant shifts point to polygenic selection from standing variation

The frequency shifts at outlier loci argue against a simple sweep. A hard sweep strips diversity around the target as one haplotype climbs toward fixation. We observed the opposite: expected heterozygosity (2*pq*) rose at selected outliers relative to matched control loci. This pattern arises when variants that are rarer at the beginning of the experiment rise to intermediate frequency (i.e., become more common) by the end of the experiment. This pattern is consistent with polygenic adaptation from standing variation with subtle shifts in allele frequency [68,75]. Standing variation, including cryptic variation released in a new environment, is the expected substrate for rapid larval adaptation of this kind [25].

Our results are in line with other larval selection experiments with regard to selection on standing variation. Brennan et al. (2019) found soft sweeps on standing variation through elevated linkage disequilibrium, although in contrast to our study they found that overall diversity declined under selection [22]. This pattern may result from selection on common variants at the beginning of the experiment that rise closer to fixation by the end of the experiment. Similarly, Bitter et al. (2019) saw phenotypic variation widen under low pH, which they read as cryptic variation exposed in a new environment [21]. We did not test for cryptic variation directly, but the Private tier shows a similar signature: variants rare at the start of the experiment, with no detectable frequency change in controls, that rose only under stress.

### Limitations

Although we took steps to control for confounding factors (e.g., by including a control treatment and removing loci from analysis that changed frequency through time in the control), there are a few important caveats to our experiment. Selection ran over a single 24-hour window of early larval development, and sensitivity to acidification and effluent probably shifts across later larval and juvenile stages, where carryover effects on energy budgets and settlement are well documented in bivalves [14]. Pooled exome capture gives allele frequencies, not haplotypes or genotypes, so we cannot separate hard from soft sweeps cleanly. Capture efficiency can also differ among individuals within a pool, which would bias pooled allele-frequency estimates; the control filter removes only biases shared across treatments, so some residual capture bias may remain. We interpret the stressor interaction from the control-anchored contrast rather than a single genome-wide interaction estimate, because the per-locus interaction did not repeat across spawns and one summary number would imply a precision the data do not support.

### Conclusions

The combined CASE response was additive in the size of allele-frequency change and synergistic in composition: it recruited loci, genes, and functions that neither single stressor did. That distinction has two implications. Single-stressor experiments will miss targets that appear only under combined exposure, and outlier loci from one genetic background may not transfer to another even when the same functions are selected, so functional categories may travel better than individual markers for predicting multistressor responses.

Coastal oyster populations encounter acidification and effluent together. Across independent spawns, enriched functions recurred more often than expected while most individual loci did not. This is important for both aquaculture and management. Our results argue for protecting rare standing variation and the large populations that maintain that variation. The rare variants that carried the largest shifts under CASE are the class most easily lost to drift, bottlenecks, and hatchery founder effects, which reinforces existing calls to conserve the large populations that retain them [21,22]. For aquaculture, our data indicate that the variants favored by mass selection will depend on the genetic background of the broodstock, even when the traits and functions selected are the same.

## Materials and methods

### Broodstock collection and conditioning

Wild adult eastern oysters (*Crassostrea virginica*) were collected under Massachusetts Division of Marine Fisheries scientific permit number 173000. We collected 50 wild oysters from Ipswich, Massachusetts across 2016 and 2017, and 240 putatively wild oysters from Barnstable, Massachusetts (60 in 2016 and 180 in 2017). Broodstock were held at 18 to 20 °C for four to eight weeks to ripen gametes and fed a commercial microalgal diet (Shellfish Diet 1800, Reed Mariculture) at roughly 2 to 4% of mean dry body weight per day.

We carried out a series of thermally induced spawns over 2016 and 2017 and used a spawn in the experiment only when both male and female contributors from Ipswich were present, which held the source population consistent across spawns. Four spawns met this criterion and became the experimental blocks: SP1 (30 August 2016), SP2 (13 July 2017), SP3 (24 July 2017), and SP4 (27 July 2017). Spawns are numbered sequentially in the order they were used; the corresponding internal laboratory block identifiers, retained in the archived data files, sample names, and sequence records, are B6, B10, B11, and B12, respectively (Table S1). Each spawn used a separate set of wild broodstock, so the spawns are independent genetic backgrounds rather than replicate cultures of one cross. SP2, SP3, and SP4 were sequenced; SP1 was phenotyped but not sequenced.

### Spawning and fertilization

Spawning was induced thermally. Conditioned adults were acclimated and then warmed gradually to 30 °C while watched for gamete release, and spawning males and females were separated into individual containers as sex was determined. Each spawn pooled gametes from multiple males and females, with Ipswich contributors present for both sexes by design (SP1: 9 males and 4 to 5 females, all from Ipswich; SP2: 3 males and 2 females; SP3: 11 males and 7 females; SP4: 9 males and 11 females; counts pool Ipswich and Barnstable contributors). Eggs and sperm were diluted in filtered seawater at 20 °C, egg density was estimated on a Sedgewick rafter slide, and sperm was added to the egg suspension. Fertilization was checked by scoring polar-body formation in a subsample, and more sperm was added if fertilization was below 90%. Fertilized embryos were passed through a 20 µm screen, rinsed into filtered seawater at 20 °C, adjusted to a density of 1,500 to 3,000 larvae per mL, and held at 20 °C for roughly 15 hours until the start of exposure (early larvae, trochophore stage).

### Multiple-stressor exposure experiment

Larvae from each spawn were exposed in a 2 x 2 factorial design crossing coastal acidification (CA) and sewage effluent (SE): control (CON), CA alone, SE alone, and the combination (CASE). Each treatment was assigned to three replicate 1 L glass bottles, for 12 bottles per spawn, and treatments were assigned to bottles at random. Larvae were stocked at approximately 85 per mL in 1,050 mL for SP1 (about 93,500 larvae per bottle) and at roughly 118,000 to 120,000 larvae per bottle for SP2 through SP4. During SP2, two bottles, (CA J05 and CA 08) mistakenly had the wrong treatment water added at first. Both bottles were immediately wash and restarted. We also added two additional CA bottles (that skipped the first acclimation). For analysis, we replaced J05 with one of the additional bottles to mitigate any effects of the restart or additional bottles that skipped acclimation.

Treatment water was prepared on a gas-mixing system (Ries Lab) that bubbled filtered seawater to set points of approximately 400 ppm CO_2_ (control) and 2,800 ppm CO_2_ (acidified). Sewage effluent was collected from a large urban east coast publicly owned treatment works using secondary treatment on the morning of each experiment, returned to the laboratory, and bubbled to the matching CO_2_ level (control or acidified) for at least 12 hours before use. SE treatments received 5% effluent by volume (a mix of 7.5 L treatment seawater with 0.395 L effluent); non-SE treatments received the same volume of CO_2_-equilibrated deionized water in place of effluent. Larvae were exposed for 24 hours in capped bottles at approximately 19 °C and a salinity of approximately 29.

### Larval survival and morphometrics

Bottles were subsampled at the start of exposure (T0) and at roughly 12 and 24 hours. At each point, three 1 mL aliquots were drawn from each bottle, stained with neutral red, and fixed in 10% buffered formalin. Larvae were counted on a Sedgewick rafter slide and classified as normal, deformed, or dead. Percent survival at 24 hours was calculated relative to the T0 count for each bottle. For morphometrics, at least 30 normal larvae per sample (plus any abnormal larvae) were photographed on a compound microscope (OMAX) at 100x with ToupView, and shell height, hinge length, and circumference were measured in ImageJ [76], with the scale set against the Sedgewick rafter grid. Larvae were staged as trochophore, D-stage veliger, or veliger. Survival was expressed relative to a composite initial (T0) density: the mean of the stocking (seeded) density, the jar’s own calculated T0 density sample, and the spawn-mean calculated T0 density. This composite denominator mitigates the sampling error introduced by under-homogenizing near-full culture containers when taking initial density samples; this composite denominator was used for all survival statistics, and per-jar survival under each individual denominator is displayed in Fig S2.

Survival was analyzed on spawn-centered composite survival (survival minus the spawn mean, to remove among-spawn baseline differences) with a one-way analysis of variance followed by Tukey honestly significant difference tests, and the CA × SE interaction was tested with a factorial betabinomial generalized linear mixed model (glmmTMB) using the composite T0 larval count as the binomial denominator and a random spawn effect; an ordered-beta model on the survival proportion gave the same result. The additive expectation for CASE survival was computed from the separate CA and SE effects relative to the control and compared to observed CASE survival to test for phenotype synergy.

### Carbonate chemistry and water quality

Carbonate chemistry was measured at two points: an initial sample taken once per treatment from the mixing tank in which that treatment was made, and a post-exposure sample taken from each of the 12 bottles at the end of the exposure. Temperature, salinity, and pH (total scale, pHT) were measured with a glass thermometer (precision 0.1 °C), a YSI 3200 conductivity probe (0.1 ppt), and an Accumet solid-state pH electrode (1 mV) calibrated with pH 7.01 and 10.01 NBS buffers (for slope) and Dickson seawater Certified Reference Material (CRM, for intercept) to account for the effect of salinity on pH. For total alkalinity (TA) and dissolved inorganic carbon (DIC), seawater was collected in 250 mL borosilicate glass bottles sealed with a vacuum-greased ground-glass stopper, poisoned with 100 µl of saturated mercuric chloride solution, and refrigerated at 6 °C until analysis. TA and DIC were measured on a VINDTA 3C system (Marianda Corporation): TA by open-cell potentiometric Gran titration and DIC by coulometry (UIC 5014), both with a precision of about 3 µmol kg-seawater-1 determined by repeated measurement of Dickson CRM. The full carbonate system (pHT, pCO_2_, calcite and aragonite saturation states, and carbonate, bicarbonate, and aqueous CO_2_ concentrations) was calculated from salinity, temperature, DIC, and TA with CO2SYS version 2.1 [77], using carbonic-acid dissociation constants (K1, K2) from Roy et al. [78], the hydrogen-sulfate constant (KHSO4) from Dickson [79], and the boron concentration from Lee et al. [80]. Both TA and DIC were measured directly for every bottle in this experiment, so no substitution or estimation was required.

### Larval collection for sequencing

At the start of exposure, four pools of roughly 40,000 larvae each were filtered from the starting larval stock and flash-frozen as initial (T0) samples. At the end of the exposure, larvae from each bottle were concentrated on a stack of 20, 40, and 70 µm cell strainers, rinsed into filtered seawater, transferred to filter inserts, and preserved (RNAlater, then liquid nitrogen). Survival and carbonate chemistry were recorded on a common set of twelve bottles per spawn (three per treatment). In SP3 and SP4, larval yield was sufficient to set up additional bottles beyond this set; these extra bottles were used for genomic sampling only and were not scored for survival or sampled for carbonate chemistry. For sequencing, therefore, each treatment within a spawn was represented by replicate post-exposure pools, each paired with an initial pool: three initial- and-post replicate pairs per treatment in SP2, and four per treatment in SP3 and SP4.

### DNA extraction and EecSeq library preparation

Genomic DNA and total RNA were co-extracted from each larval pool with the Zymo Quick-DNA/RNA Microprep Plus Kit, providing the genomic DNA for capture libraries and the RNA for probe synthesis from the same samples. Genomic DNA was sheared to approximately 150 bp on a Qsonica sonicator (27 minutes total, 15 s on and 15 s off, 25% amplitude), and genomic libraries were built from 500 ng of sheared DNA with the KAPA HyperPrep Kit (end repair and A-tailing, adapter ligation, and indexed amplification with dual i5 and i7 index primers).

Capture probes were synthesized from the experimental larvae following the EecSeq protocol [30]. For each spawn, stranded mRNA libraries were prepared from 1 µg of total RNA from the 24 h larval pools with the KAPA Stranded mRNA-Seq Kit (half reactions). These cDNA libraries were pooled at 200 ng each, normalized by duplex-specific nuclease (DSN) treatment, and reamplified. The sequencing adapters were then cleaved by restriction digestion using SA1I-HF (NEB) and removed with mung bean nuclease, and the remaining exon-derived fragments were biotinylated by Klenow extension to make the spawn-specific probe set.

For capture, 200 ng of each genomic DNA library was combined with 500 ng of biotinylated probes and hybridized for 48 hours at 65 °C in a buffer containing SSC, Denhardt’s solution, EDTA, SDS, human COT-1 DNA, and blocking oligonucleotides. Probe-bound fragments were captured on streptavidin beads (Dynabeads M-280), washed through a decreasing-stringency SSC series, eluted at 80 °C, and amplified by a 12-cycle post-capture PCR. Captured libraries were sequenced on the Illumina platform in 150 bp paired-end mode. The 57 libraries carried into analysis yielded 6.6 to 31.0 million read pairs each (median 16.2 million), with a median primary mapping rate of 98.1% (range 82.4 to 98.8%).

### Read processing, mapping, and variant calling

Reads were quality trimmed with fastp [81] using sliding-window trimming from both ends (window size 5, mean quality 15), a per-base quality threshold of Q15, removal of reads with more than 50% low-quality bases, paired-end adapter auto-detection and overlap correction, and a minimum retained length of half the input read length. Trimmed reads were mapped to the *C. virginica* reference genome (C_virginica-3.0; chromosomes 1 to 10 correspond to RefSeq accessions NC_035780.1 to NC_035789.1) [31,32] with BWA-MEM [82] using a match score of 1, a mismatch penalty of 4, and a gap-opening penalty of 6, marking shorter split hits as secondary, through the dDocent pipeline [83] (script dDocent_ngs_3.1). Duplicates were marked with Picard MarkDuplicates (Broad Institute: https://broadinstitute.github.io/picard/; optical-duplicate pixel distance 2,500), and alignments were filtered to mapping quality of at least 10, excluding secondary and duplicate alignments and heavily soft-clipped reads. Variants were called with FreeBayes [84] in pooled-continuous mode (minimum mapping quality 5, minimum base quality 5, minimum alternate fraction 0.025, minimum alternate count 1, up to 20 alleles per site). The raw call set was reduced with VCFtools [85] to sites genotyped in at least 90% of pools with a non-reference allele frequency between 0.001 and 0.9999, a minor allele count of at least 1, and a minimum quality of 30, then exported for project-specific filtering.

### Variant filtering and pooled allele-frequency estimation

Variants were filtered with a custom pipeline (CASE_PVCF_filter2.sh). Within each spawn, we retained variants (loci) with less than 10% missing data, less than four alleles, and a minor allele frequency above 0.015 (alternate allele frequency between 0.015 and 0.985), computed from read counts with bcftools. Loci passing these filters in all three sequenced spawns were combined into one synchronized file with VCFtoPopPool.py and popoolation2 [86], and we required a per-pool minimum coverage of 10 and a total coverage above 24. Pooled allele frequencies were estimated from the sync files with the poolSeq package (read.sync and af, rising polarization) [74].

Our experimental design had four T0 samples of larvae from the initial larval pool with single larval pools from replicate exposure bottles per treatment. The bottles are independent, but the initial larval samples are not. To help create coverage balanced initial samples, we pooled all read counts across the four initial samples, and then created three or four random draws (depending on the spawn)

### Detection of selected loci: CMH tests and FDR control

Within each spawn, larvae were sampled from a common pool prior to treatment, so the four initial-sample (IS) libraries represent replicate sequencing of a single pre-exposure population rather than independent experimental units. For each spawn, allele counts from the four IS libraries were therefore summed site-by-site into a single T0 pool (`sum.sh`), and that pool was replicated to match the number of post-exposure treatment replicates in that spawn (three in SP2; four in SP3 and SP4). To ensure that pre- and post-exposure pools were comparable in sequencing depth, we calculated the genome-wide mean per-pool coverage across all post-exposure treatment libraries (AV_COV) and retained only sites at which the summed IS depth met or exceeded this value; per-pool treatment depths were then capped at the corresponding IS depth. Each T0 pseudo-replicate was then generated independently by randomly resampling alleles at each site from the summed IS counts (`sub_sample.py`), with a target depth equal to the summed IS depth divided by four, the average depth of a single IS library. Draws used a fixed random seed, offset deterministically by site, so that the resampling is exactly reproducible. This procedure yielded, for each spawn and each treatment (CA, SE, CASE, CON), a paired matrix of T0 and post-exposure allele counts that was depth-matched across timepoints and in which each replicate carried its own independent sampling variance; these matrices were supplied to the adapted CMH test as generation 0 and generation 1, respectively.

For each treatment within each spawn, we tested for consistent allele-frequency change between the initial and post-exposure pools across replicates with the adapted Cochran-Mantel-Haenszel (CMH) test in the ACER package [73], which accounts for genetic drift and the extra sampling noise of pooled sequencing. Effective population size was set to 250 for SP2 (due to the lower number of breeders) and 1,000 for SP3 and SP4. Pool sizes were 40,000 for the initial pools and 10,000 for the post-exposure pools (the conservative lower bound of our estimates; see Results), the starting frequency was the replicate mean, and a minimum coverage of 20 (SP2 and SP4) or 14 (SP3) was applied. The same test was run on all three spawns combined (the across-spawns analysis). Raw p-values were converted to q-values with the qvalue package [87] using pi0 = 1, which is equivalent to a Benjamini-Hochberg false discovery rate.

### Locus tier assignment

Significant loci were assigned hierarchically to one of three tiers (Core, then Convergent, then Private), so each locus appears once. Core loci show replicated, same-direction signal across independent spawns: Tier A loci were significant at q < 0.10 in all three spawns, and Tier B loci at q < 0.05 in at least two spawns. Convergent loci show weak but consistent signal detectable only when spawns are pooled (per-spawn q < 0.01 in at least one spawn and across-spawns q < 0.0001). Private loci are strong single-spawn outliers among loci testable in only one or two spawns, taken as the top 1% of treatment q-values within the significant set using spawn-specific thresholds (SP2 q < 0.001, SP3 q < 0.1, SP4 q < 0.01) to account for differences in drift and power among spawns. Low-variation singletons whose pooled signal was driven by a single polymorphic spawn were reclassified here from the Convergent candidates.

### Control-treatment filtering

To reduce experimental artifacts and family effects, loci that also changed in the control were removed. Loci with a control q-value below 0.10 were excluded from the treatment sets, and loci overlapping any gene region that contained a control-significant locus were removed with bedtools [88]. Significant-locus intervals were defined by merging locus positions with a 100 bp flank before annotation.

### Gene annotation and Gene Ontology enrichment

Significant loci were annotated to genes by intersecting their positions with the *C. virginica* gene model (bedtools intersect against sorted.ref3.0.gene.bed) and recording NCBI LOC gene identifiers. Gene Ontology (GO) enrichment was tested for each stressor (CA, SE, CASE), for all outliers combined, and for each locus tier with topGO [89] using Fisher’s exact test. The gene-to-GO mapping was built from a Gene Ontology annotation file (GO/gene_ontology.gaf) derived from the NCBI RefSeq annotation of the *C. virginica* genome assembly GCF_002022765.2 (C_virginica-3.0).

To summarize function without imposing predefined categories, the enriched GO terms (pooled across stressors) were grouped into data-driven clusters by the overlap of the candidate genes annotated to each term (Jaccard index of at least 0.10, terms with at least two genes), using greedy-modularity community detection on the term-similarity network [90,91]. Clusters with at least three terms were retained and named from their member terms, and a set of stress-relevant clusters was shown in the main mechanism figure (Fig 4D) and summarized in Table 1; all clusters, including growth and developmental categories expected for early larvae, are listed in Table S7 with cluster IDs shared between the two tables. The clustering is data-driven and reproducible; the stress designation and cluster names are an interpretive overlay.

### Cross-spawn reproducibility analysis

To test whether selection was reproducible across independent spawns, we counted treatment-outlier loci significant in all three spawns (per-spawn *q* < 0.10) and in at least two (per-spawn *q* < 0.05; SP3 *q* < 0.10), matching the Core tier rules. Expected overlap came from 500 permutations that drew, for each spawn, the same number of loci at random from that spawn’s eligible loci, preserving differences in probe sets among spawns. As a baseline, we built a control outlier set by applying identical tier rules to control *q*-values. The control set was analyzed unfiltered and filtered reciprocally, excluding loci with any treatment *q* < 0.10 and loci significant in treatments both within and across spawns, mirroring the control filter applied to treatments. Treatment and control enrichments were compared as a ratio of observed-to-expected rates with a conditional binomial test.

Outlier loci from each spawn (two-spawn thresholds) were mapped to genes. Expected gene recurrence came from 500 draws of the same number of distinct genes per spawn, sampled from genes captured in that spawn with probability proportional to their number of captured loci; drawing from all annotated genes inflated recurrence for every set, including the control, because captured genes overlap among spawns. GO terms were tested for enrichment within each spawn (hypergeometric test on propagated topGO annotations, all annotated genes as background, *p* < 0.01, at least three genes), and recurrence of enriched terms across spawns was compared with 200 random gene sets drawn the same way. Treatment specificity was assessed by comparing overlap between the same treatment in two spawns with overlap between different treatments, permuting treatment labels within spawn (5,000 permutations).

The control served as a positive control for a repeatable response independent of treatment.

### Allele-frequency architecture

For each tier we summarized the starting minor allele frequency (the replicate-mean initial frequency) and the magnitude of allele-frequency change (the absolute difference between post-exposure and initial frequency), per treatment and spawn. These distributions were compared across tiers to characterize the frequency spectrum and effect size of each mode of response.

To test whether the combined-stressor genomic response was additive in magnitude, we computed the per-locus allele-frequency change for each treatment and spawn (mean of post-minus-initial change across replicate pre/post pool pairs, requiring per-pool coverage of at least 20x) and compared the control-anchored CASE change (ΔAF_CASE_ − ΔAF_CON_) to the additive expectation from the single stressors ((ΔAF_CA_ − ΔAF_CON_) + (ΔAF_SE_ − ΔAF_CON_)), using one point per locus averaged across the spawns in which it was scored. Additivity was summarized within significance-overlap categories by the orthogonal (Deming, error-variance ratio 1) regression slope and the fraction of loci above the 1:1 line (Fig 3; by replication tier in Fig S5). Anchoring both axes on the control removes the shared handling/lab response, following the pool-seq multiple-stressor precedent [35,36].

### Diversity at selected loci

For each treatment, expected heterozygosity (*H_E_* = 2*pq*) was estimated at each locus from the pooled allele frequencies with a read-count correction of c/(c-1), where c is the per-pool coverage. At that treatment’s outlier loci, within each tier, we computed the change in He from the initial to the post-exposure pools and subtracted the change at the same loci in the control pools, yielding a control-anchored excess that removes the shared initial-to-post offset and the ascertainment that comes from selecting loci on allele-frequency change. Loci were required to have a coverage of at least 2 in both the initial and post-exposure pools. Because outlier loci are ascertained on allele-frequency change, these values index the direction and relative magnitude of diversity change at selected loci rather than genome-wide diversity. To test whether the control-anchored excess exceeds ascertainment and drift, each observed excess was compared to a frequency- and shift-matched neutral null: for each outlier locus, non-outlier loci matched on binned starting minor allele frequency and within a caliper on the magnitude of allele-frequency change were drawn (1,000 replicates), and a z-score (observed minus null mean, divided by the null standard deviation) was computed per tier and treatment; the matched-null band and z are shown in Fig 5 and the matching adequacy in Fig S14.

### Ordination

Samples were ordinated by principal component analysis of allele frequencies across three locus sets: 10,000 random genome-wide loci, the outlier loci, and each tier separately. We also ordinated the per-sample change in allele frequency relative to the initial pool. Genome-wide (Fig S3) and tier all-sample ordinations used correlation PCA (prcomp, centered and scaled); the allele-frequency-change ordination (Fig 2A) and per-spawn outlier ordinations (Fig 2A to C) used covariance PCA (centered, not scaled), and the per-spawn outlier ordinations were restricted to loci in each spawn’s coverage- and MAF-filtered position set. Variance explained by the first two components is reported on the axes.

### Statistical environment, data, and code availability

Analyses were run in R 3.6.0. Carbonate chemistry was solved in CO2SYS version 2.1. Raw sequence reads will be deposited in the NCBI Sequence Read Archive (BioProject accession to be provided upon acceptance). All analysis code, from filtered variants through every main and supplementary figure, is provided as a single reproducible R Markdown document in the project repository (https://github.com/jpuritz/Puritz_etal_CASE).

### Use of large language models

We used Claude (Anthropic; Claude Opus and Fable models, including Claude Opus 5 and 5.5 and Fable 5.0 and 5.1), accessed through the Claude Code command-line tool and the Claude desktop application between May and October 2026, for two tasks. First, for analysis code: the variant-filtering and pseudo-replication scripts and the original analysis pipeline were written by the authors before LLM use, and Claude was used to extend, refactor, comment, and debug this code; prompts were interactive natural-language requests that described the intended analysis and supplied the relevant scripts and output files. All code was reviewed and run by the authors, and key results were checked against independent reimplementations (for example, Core-tier locus counts were reproduced exactly). Second, it edited and condensed manuscript text for clarity and length, with prompts naming the target section and word limit; the authors reviewed and revised every edit. The authors take full responsibility for the accuracy and originality of the work and for the attribution of all sources.

## Acknowledgements

We thank the technicians and administrators at our partner wastewater treatment facility. Their patience and facilitation were critical for this project. We thank Justin Reis and the Reis lab for use of their probes and mass flow controllers. We thank Sara Schaal, Alan Downey-Wall, Sandi Scripa, Jaxine Wolfe, and Ryan Haldieri for assistance with running the experiment. We thank Coline Caillon, Amy Zyck, Megan Delatte-Guidry, Jacob Green, Olivia Nieves, Dalia Bercow, and Cassandra Cerasia for comments on earlier versions of this manuscript. This work was funded by RI Sea Grant (R/1822-2022-104-01; to JBP), and the College of Environmental and Life Sciences at the University of Rhode Island (JBP), USDA-NIFA Hatch Program project 1017848/RI0019-H020 (JBP). This is contribution 002 from the Puritz Lab of Marine Evolutionary Ecology.

## Supporting information

All supporting figures (Fig S1 to Fig S16) and supplementary tables (Table S1 to Table S7) are provided in the separate Supporting Information document.

## Notes

https://github.com/MarineEvoEcoLab/Puritz_etal_CASE

### Competing Interest Statement

The authors have declared no competing interest.

