## Supplemental Information for "Additive in size, synergistic in composition: selection across the eastern oyster larval genome under coastal acidification and sewage effluent"

### Supporting figures

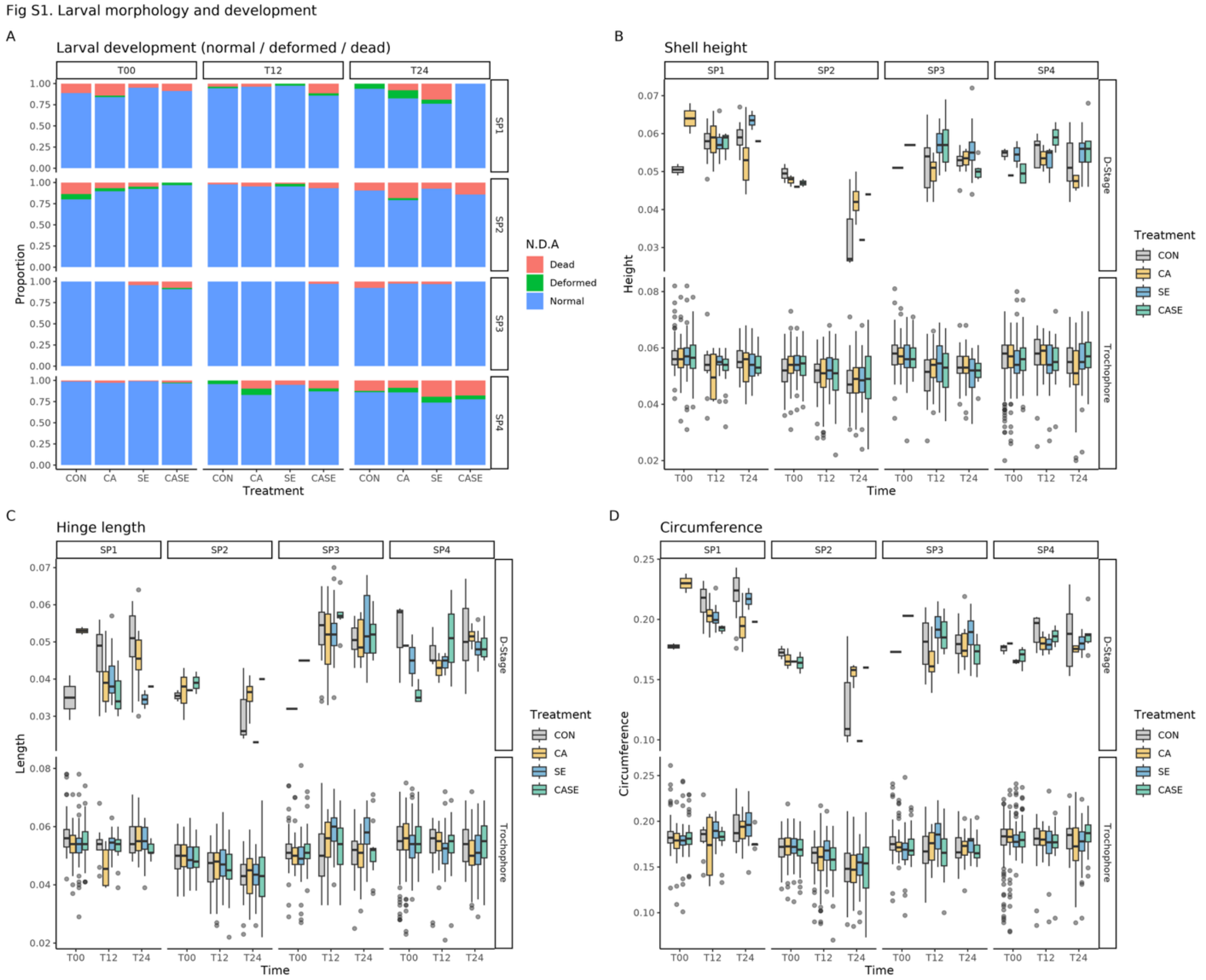


**Fig S1.** Larval morphology and development. (A) Stacked proportions of normal, deformed, and dead larvae by treatment, faceted by spawn (rows) and timepoint (columns). (B to D) Shell height (B), hinge length (C), and circumference (D) of normal, live larvae by treatment and timepoint, faceted by larval stage and spawn; box plots show the median and interquartile range.

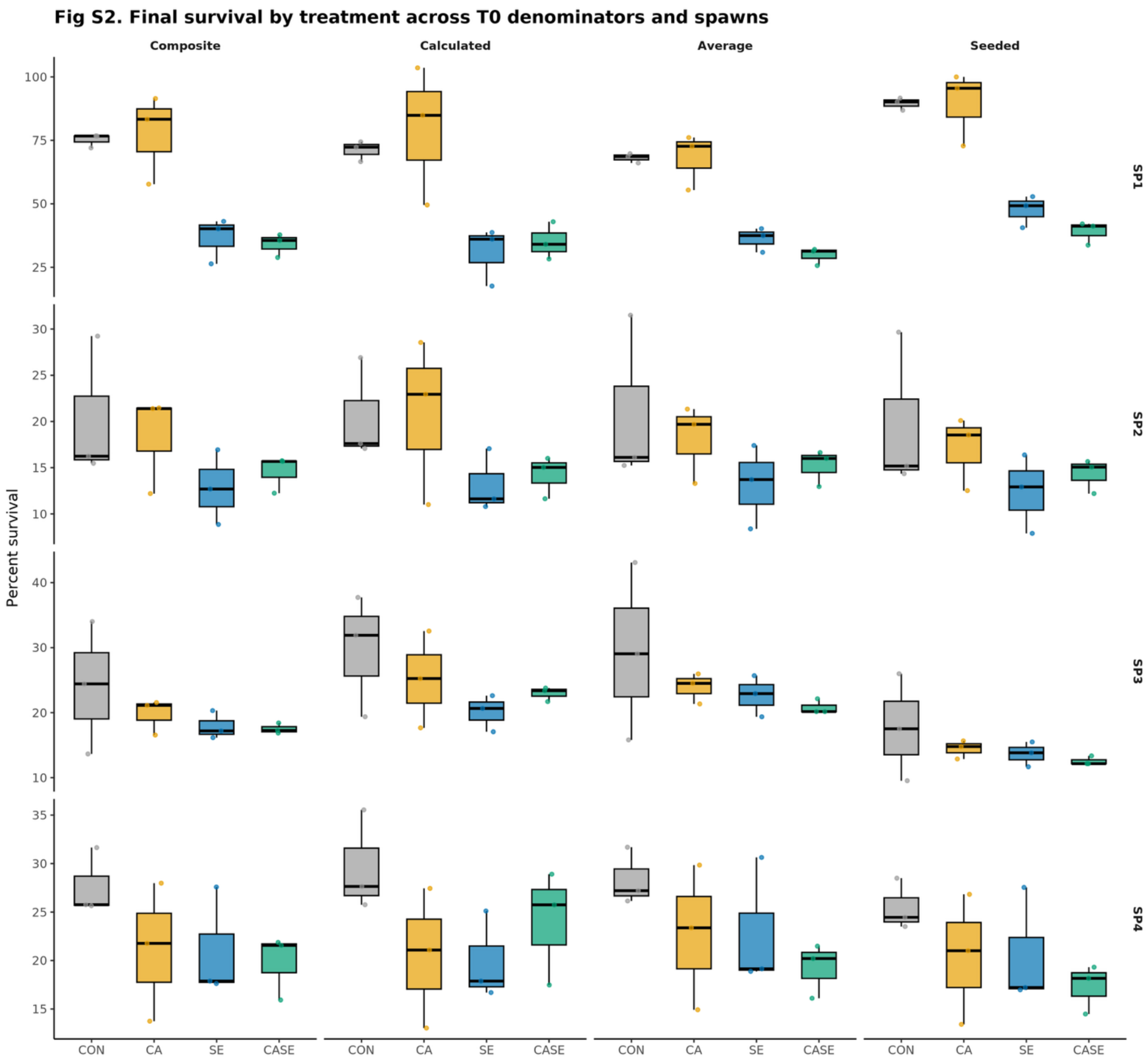


**Fig S2.** Final survival across T0 denominators. End-timepoint percent survival by treatment (CON, CA, SE, CASE), faceted by spawn (rows: SP1, SP2, SP3, SP4) and by the T0 denominator used to compute survival (columns: composite, calculated, average, seeded/stocking). Box plots show median and interquartile range with per-jar points. The treatment order (CON > CA > SE > CASE) holds across all four denominators; the composite denominator (used in Fig 1) is used for all statistics.

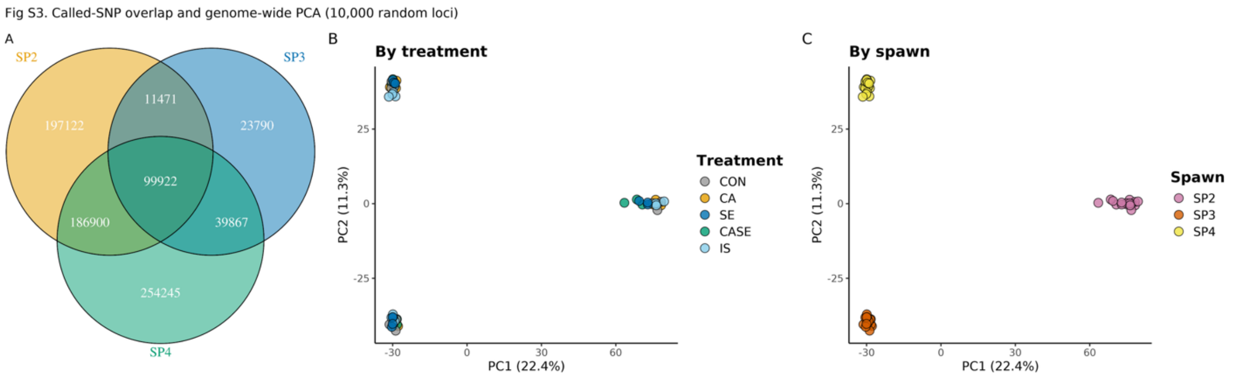


**Fig S3.** Called-locus overlap and genome-wide ordination. (A) Overlap of the loci called in each sequenced spawn (SP2, SP3, SP4). (B, C) Principal component analysis of allele frequencies at 10,000 random loci, colored by treatment (B) and by spawn (C); correlation PCA. Genome-wide structure is dominated by spawn rather than treatment, and the spawns differ in their called-locus sets, the contrast that motivates the outlier-based analyses.

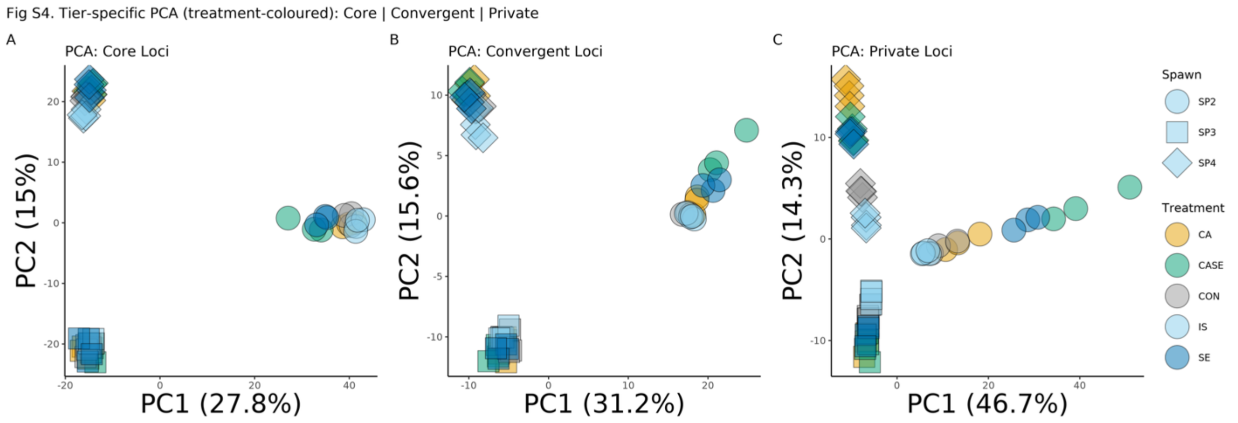


**Fig S4.** Tier-specific ordination. Principal component analysis of all samples computed separately on the Core, Convergent, and Private locus sets (correlation PCA, scaled).

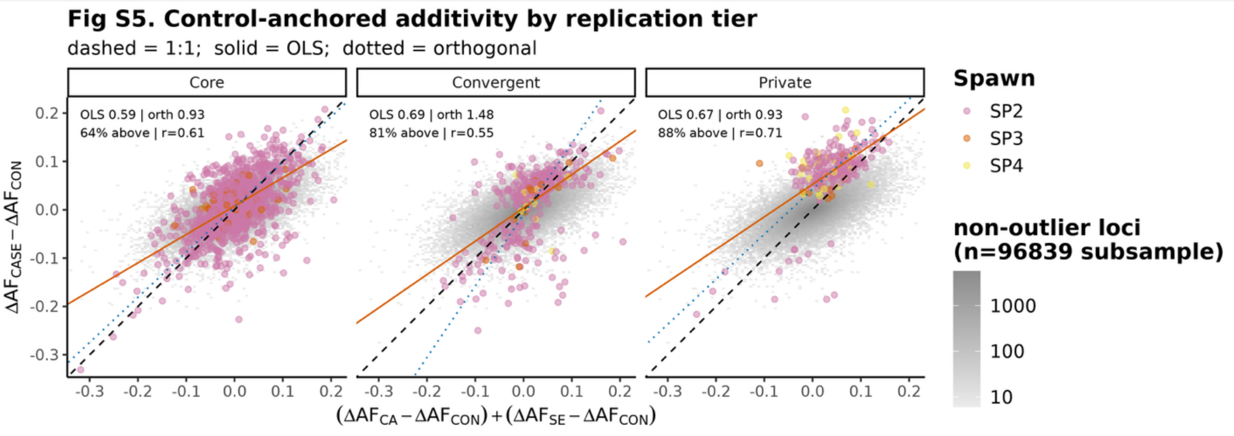


**Fig S5.** Control-anchored additivity by replication tier. The Fig 3 scatter faceted by replication tier (Core, Convergent, Private) as a robustness check, with OLS and orthogonal (Deming) slopes, percentage above the 1:1 line, and Pearson r annotated per tier; reporting OLS alongside the orthogonal fit shows the steeper orthogonal slope alone does not imply supra-additivity.

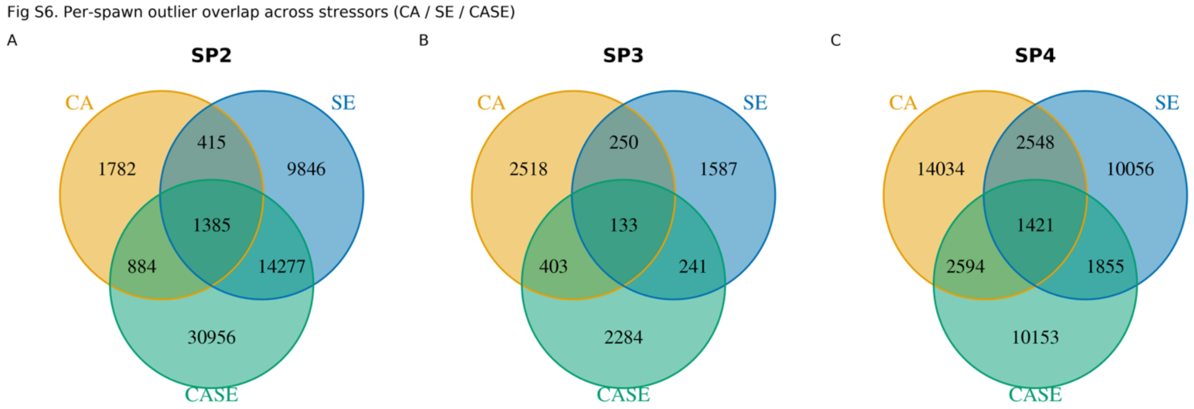


**Fig S6.** Per-spawn outlier overlap. Venn diagrams of the CA, SE, and CASE outlier loci within each sequenced spawn (SP2, SP3, SP4), shown separately, to complement the across-spawn overlap in Fig 4C.

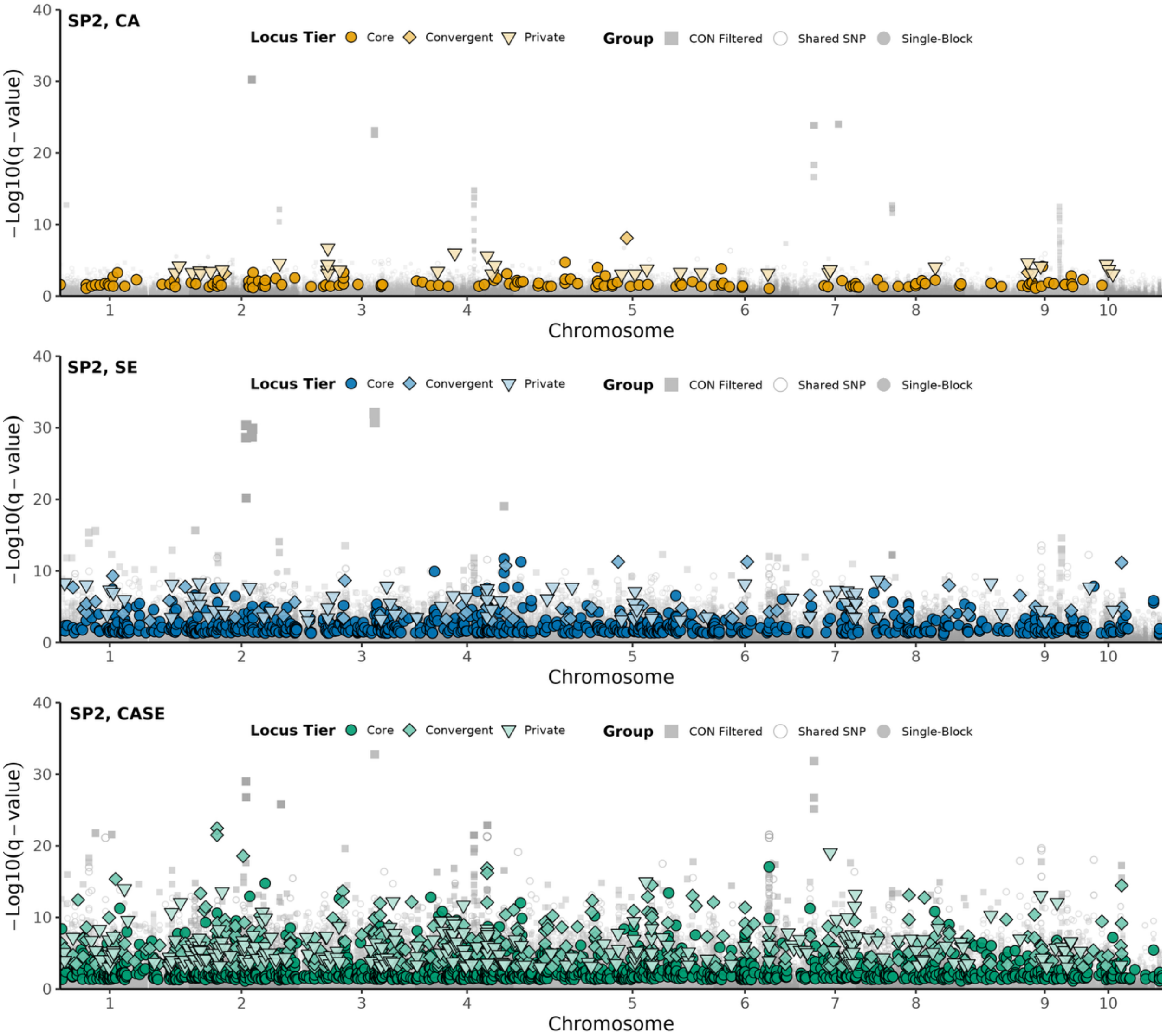


**Fig S7. Spawn SP2 Manhattan stack.** CA, SE, and CASE Manhattan plots for SP2 in the same stacked layout and tier coding as Fig 4A (y capped at 20 for cross-spawn comparability).

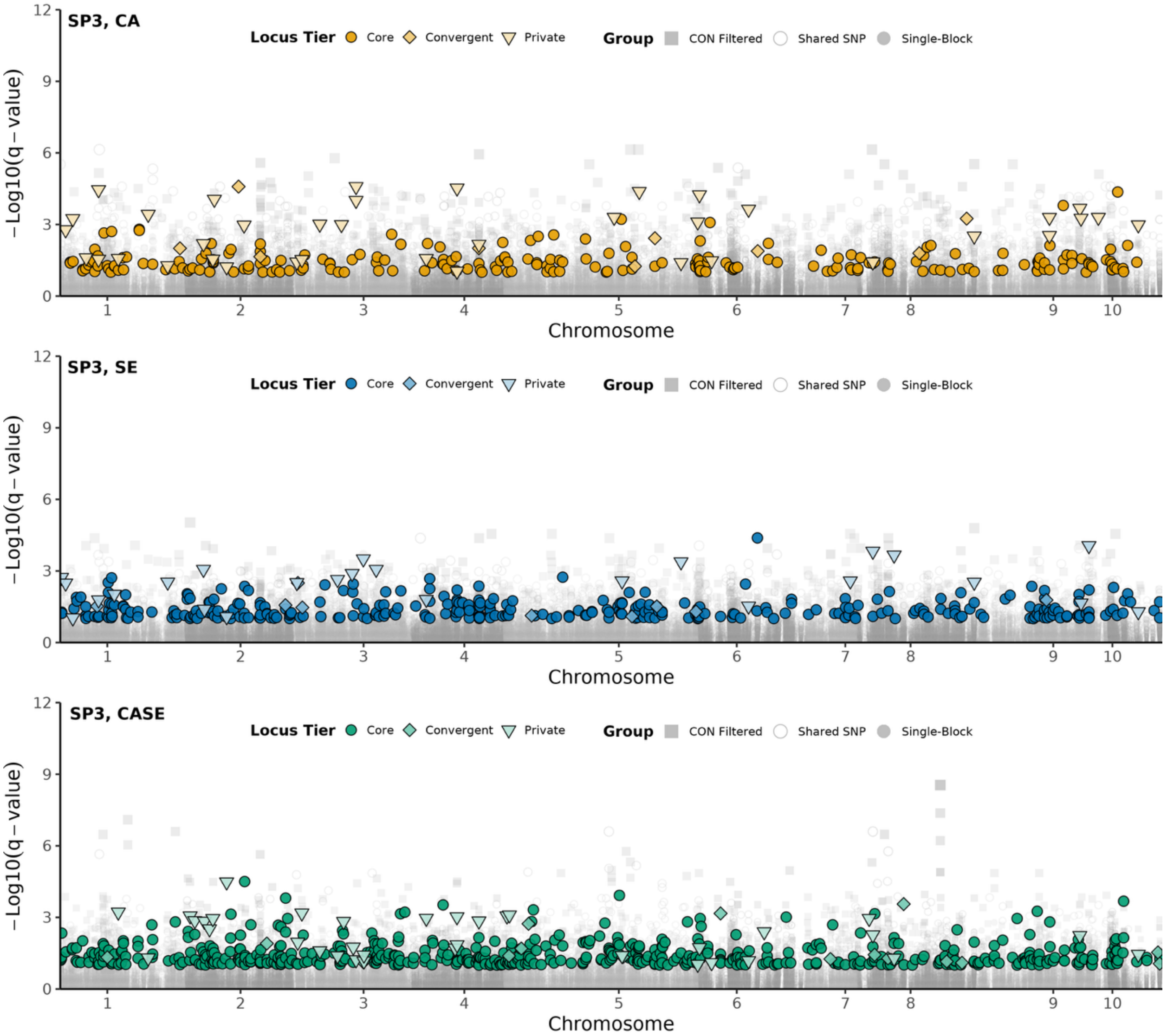


**Fig S8. Spawn SP3 Manhattan stack. As Fig S7, for SP3.**

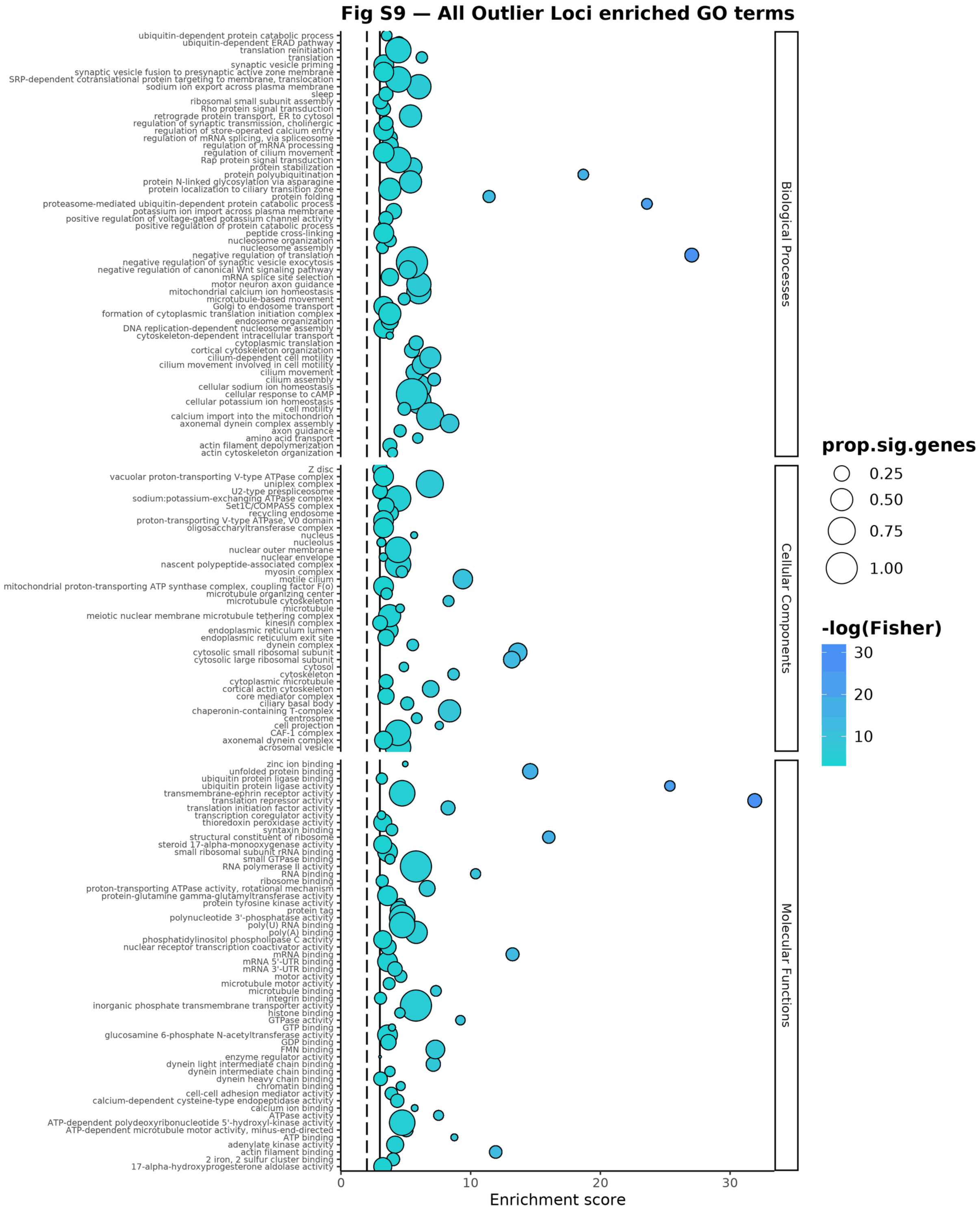


**Fig S9.** Full GO enrichment, all outlier loci. Complete topGO enrichment for the combined outlier set as a dot plot faceted by GO category; the full term set summarized in Fig 4B and tabulated in Table S5a.

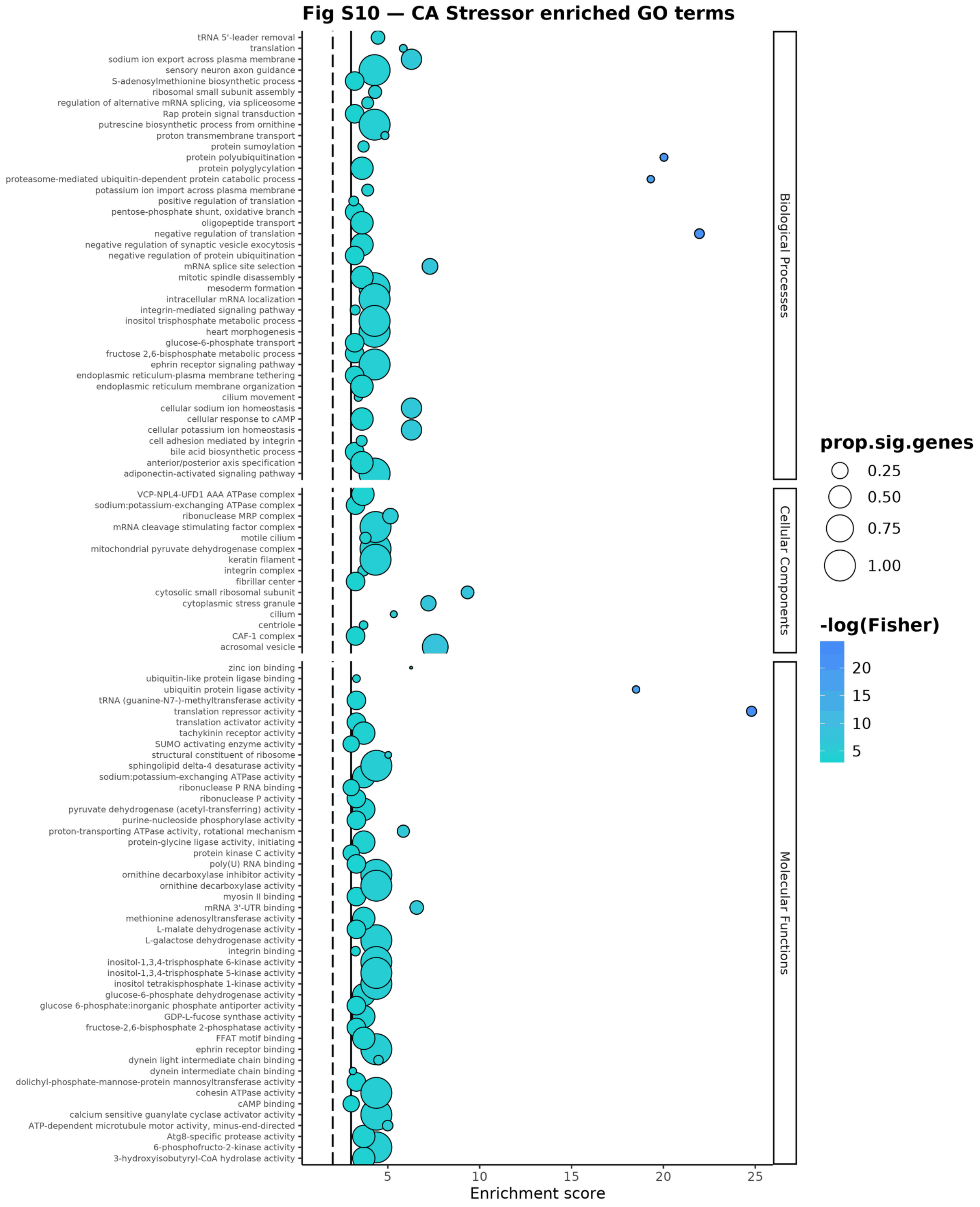


**Fig S10.** Full GO enrichment, CA. As Fig S9, for the CA outliers (Table S5b).

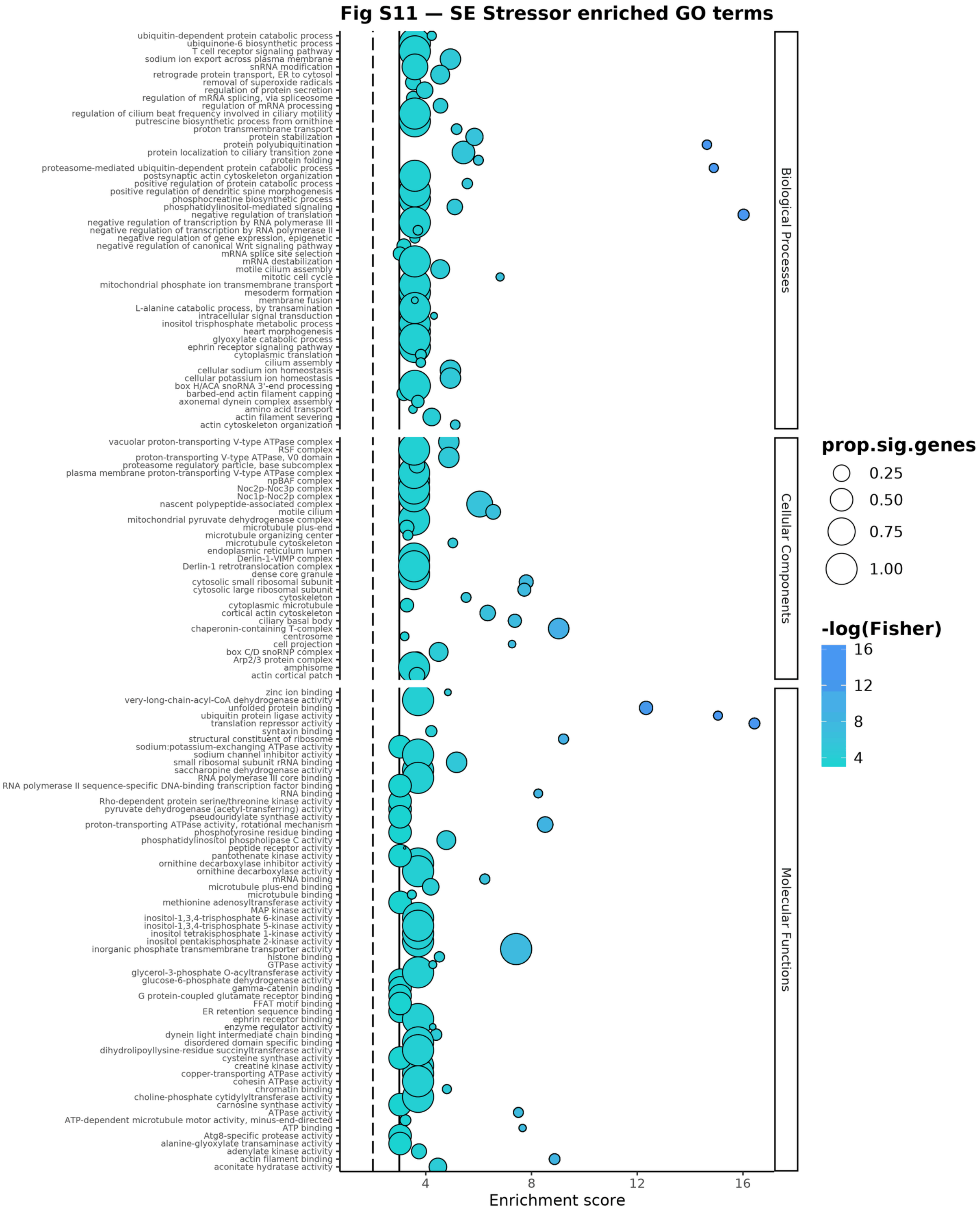


**Fig S11.** Full GO enrichment, SE. As Fig S9, for the SE outliers (Table S5c).

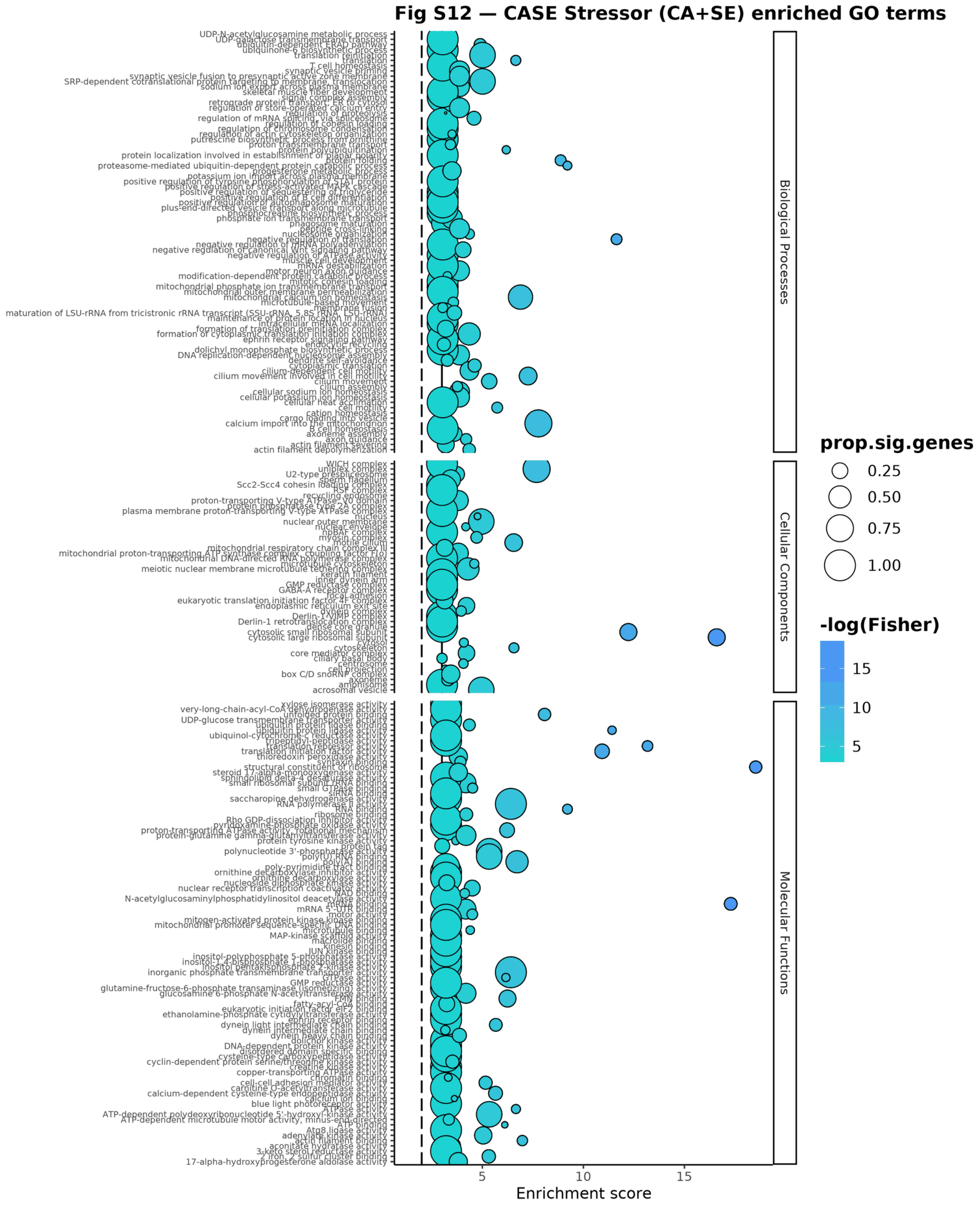


**Fig S12.** Full GO enrichment, CASE. As Fig S9, for the CASE outliers (Table S5d).

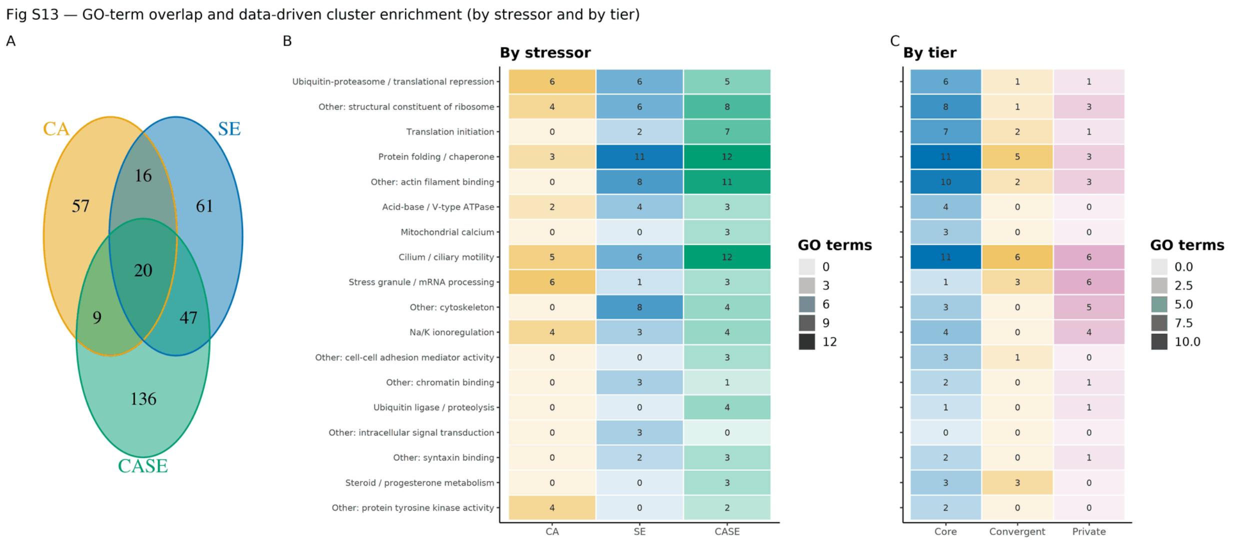


**Fig S13.** GO-term overlap and cluster enrichment. Overlap of enriched GO terms among CA, SE, and CASE, with the data-driven GO-cluster enrichment shown both by stressor and by locus tier, extending the mechanism panel in Fig 4D.

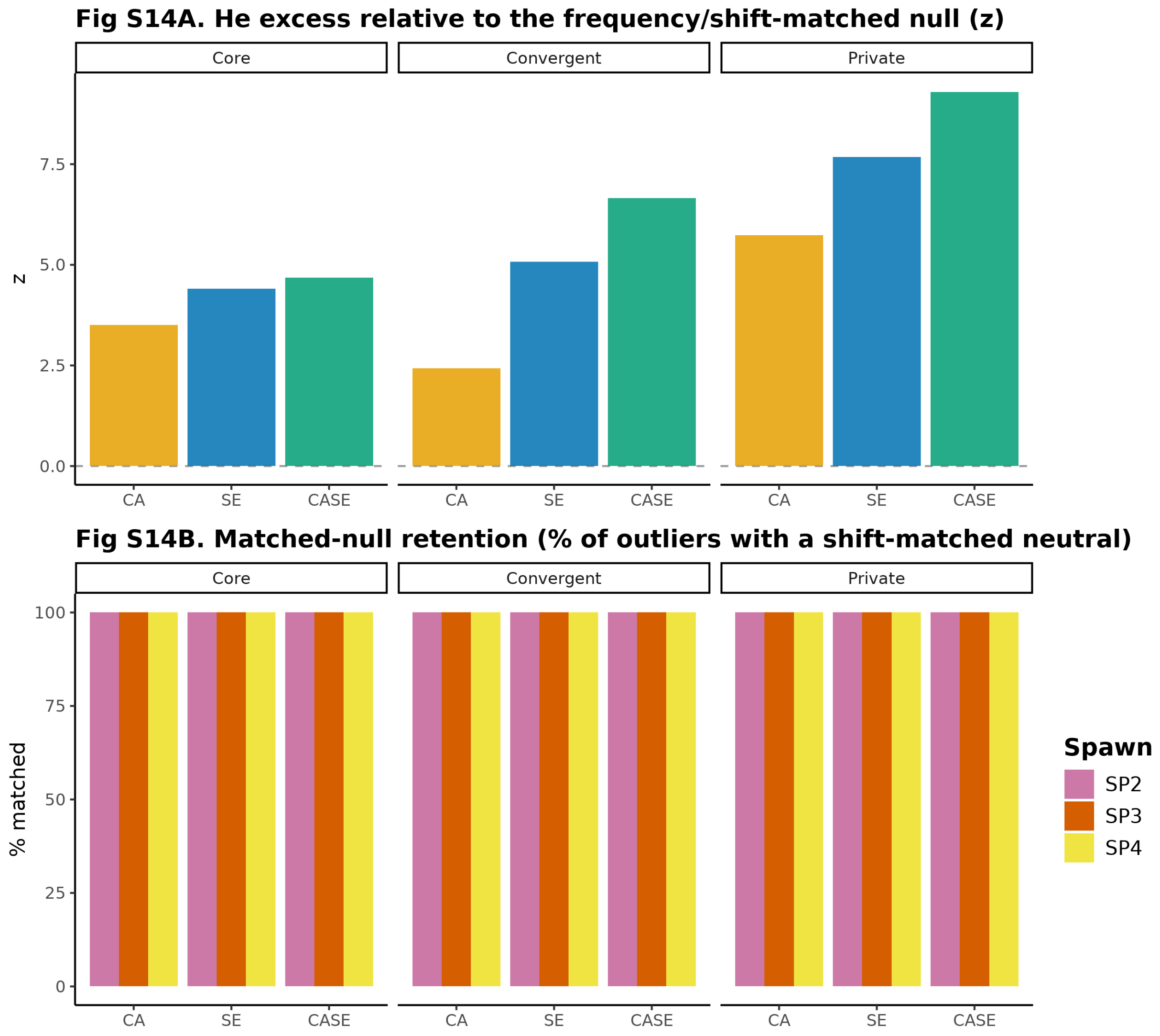


**Fig S14.** Expected-heterozygosity matched-null diagnostics. (A) The z of the observed control-anchored He excess relative to a frequency- and shift-matched neutral null, by tier and stressor (support for the null band and z in Fig 5). (B) Matching adequacy: the percentage of outliers with a shift-matched neutral locus, by tier, stressor, and spawn; interpret the z only where retention is adequate.

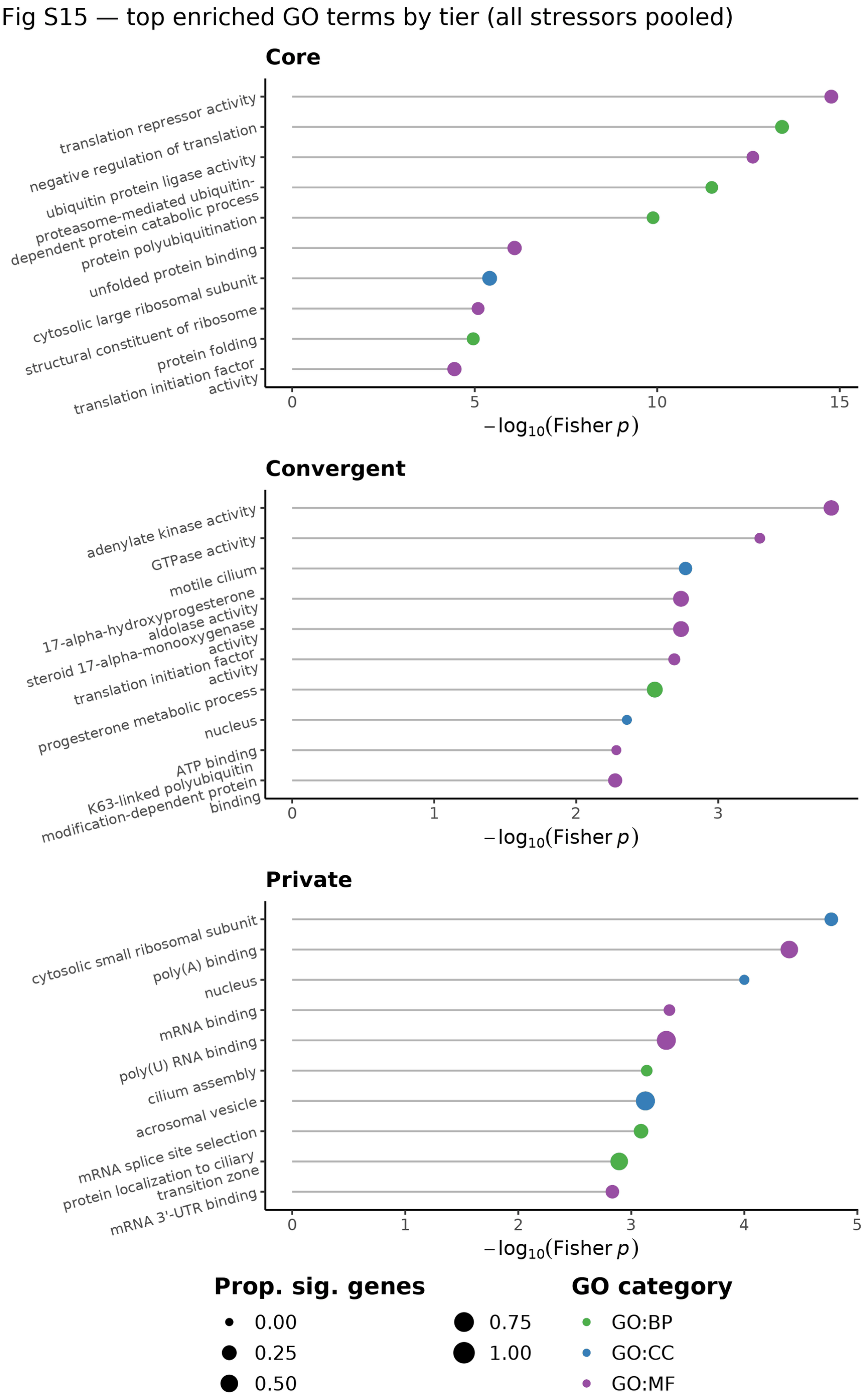


**Fig S15.** Top enriched GO terms by locus tier. Lollipop plot of the most enriched GO terms (top 10 by Fisher p, topGO) for each tier (Core, Convergent, Private), all stressors pooled; x = -log10(Fisher p), point size = proportion of significant genes annotated to the term, colour = GO category (BP, CC, MF). One panel per tier.

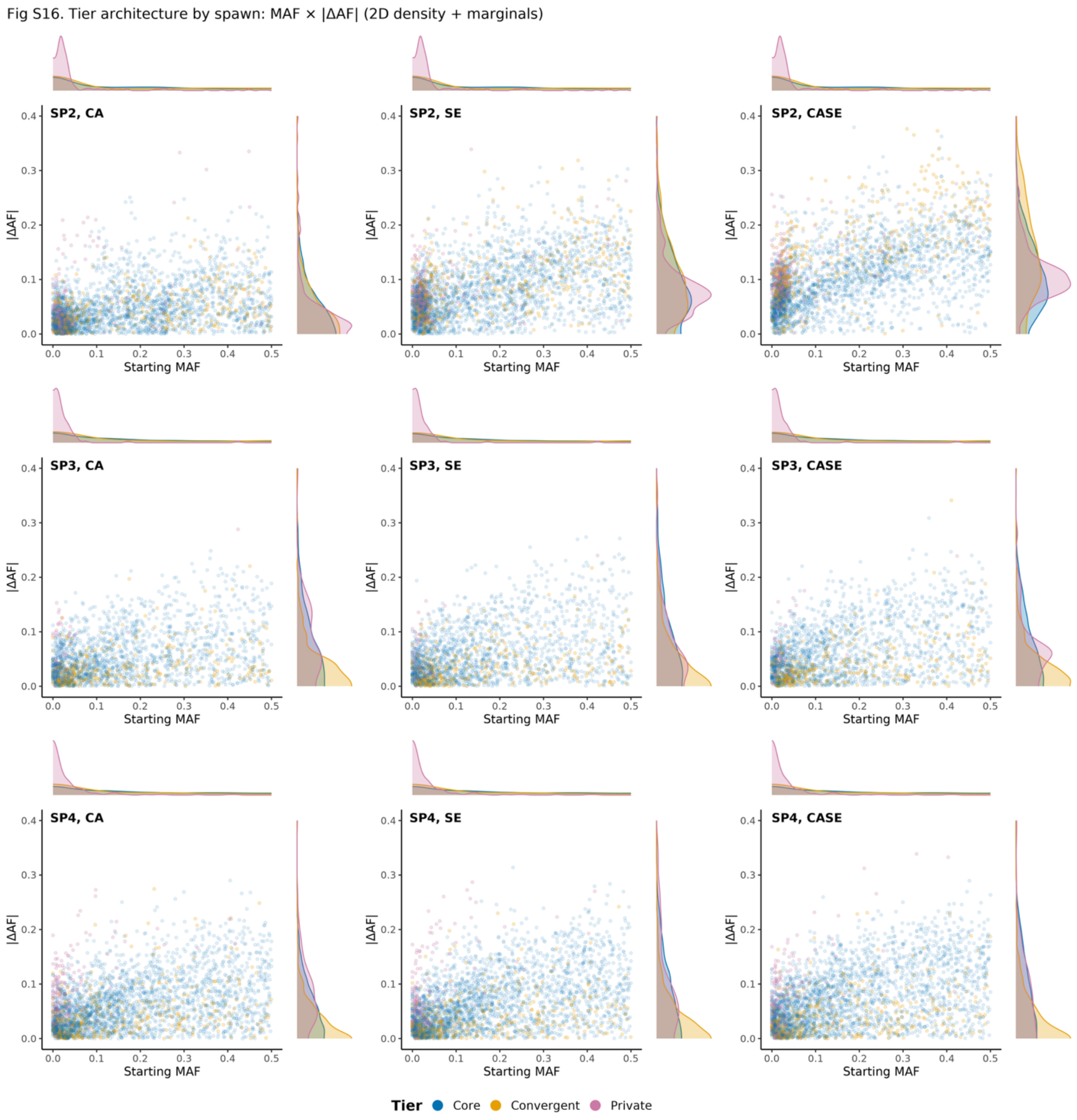


**Fig S16.** Tier architecture by spawn. The same MAF0 by |ΔAF| two-dimensional density composite as Fig 5, arranged as a 3 x 3 grid (rows = spawn SP2, SP3, SP4; columns = stressor CA, SE, CASE) with points coloured by tier, showing the tier architecture is consistent across spawns.

### Supplementary tables

Table S1. Spawn identifier mapping. Sequential spawn labels used throughout the manuscript (SP1-SP4), with spawn date, the internal laboratory block identifier retained in the archived data files, sample names, and sequence records, and the data collected for each spawn. Provided as a machine-readable file in the project repository: analysis/results/tables/TableS1_spawn_id_mapping.csv.

Table S2. Carbonate chemistry. Salinity, temperature, total alkalinity, dissolved inorganic carbon, pH (total scale), $pCO_2$, and aragonite saturation state by spawn, treatment, and sampling phase (initial mixing-tank sample; post-exposure bottles), given as mean ± SD where $n > 1$. Carbonate parameters are reported at the measured in-situ temperature (~19 °C). Both TA and DIC were measured directly for every sample and the full system solved in CO2SYS v2.1. Provided as a machine-readable file in the project repository: analysis/results/tables/TableS2_carbonate_chemistry.csv.

Table S3. Locus counts by tier and treatment, with tier totals. Counts are unique loci; a locus significant for more than one treatment is counted under each of those treatments but once in the tier total.

Table S4. Full significant-locus table with tier assignment and per-treatment significance flags. Provided as a machine-readable file in the project repository: analysis/results/tables/TableS4_significant_loci.csv.

Table S5. GO enrichment (topGO), with readable per-set tables for all outliers (S5a), CA (S5b), SE (S5c), and CASE (S5d), one row per term sorted by Fisher p. Provided as a machine-readable file in the project repository: analysis/results/tables/TableS5_GO_enrichment.csv.

Table S6. Candidate gene (LOC) lists per stressor. Provided as a machine-readable file in the project repository: analysis/results/tables/TableS6_candidate_genes_{CA,SE,CASE}.txt.

Table S7. Data-driven GO clusters. The table below summarizes every retained cluster (ID, cluster name, stress-relevant flag, CASE-only flag, number of enriched terms, number of member genes); cluster IDs are shared with Table 1 in the main text. The full member-gene listing, one row per cluster, term, and gene with the enriched stressor sets flagged, is provided as a machine-readable file in the project repository: analysis/results/tables/TableS7_GO_cluster_member_genes.csv (cluster summary: TableS7_cluster_summary.csv).

| **Tier** | **CA** | **SE** | **CASE** | **Total** |
|---|---|---|---|---|
| Core | 326 | 822 | 1479 | 2436 |
| Convergent | 31 | 83 | 406 | 493 |
| Private | 178 | 175 | 370 | 543 |

| **ID** | **Cluster** | **Stress-relevant** | **CASE-only** | **Terms** | **Member genes** |
|---|---|---|---|---|---|
| 1 | Ubiquitin-proteasome / translational repression | yes | no | 6 | 139 |
| 2 | Other: structural constituent of ribosome | no | no | 9 | 66 |
| 3 | Translation initiation | yes | no | 7 | 28 |
| 4 | Protein folding / chaperone | yes | no | 15 | 133 |
| 5 | Other: actin filament binding | no | no | 16 | 36 |
| 6 | Acid-base / V-type ATPase | yes | no | 4 | 10 |
| 7 | Mitochondrial calcium | yes | yes | 3 | 3 |
| 8 | Cilium / ciliary motility | yes | no | 16 | 44 |
| 9 | Stress granule / mRNA processing | yes | no | 8 | 11 |
| 10 | Other: cytoskeleton | no | no | 10 | 29 |
| 11 | Na/K ionoregulation | yes | no | 4 | 4 |
| 12 | Other: cell-cell adhesion mediator activity | no | yes | 3 | 10 |
| 13 | Other: chromatin binding | no | no | 3 | 16 |
| 14 | Ubiquitin ligase / proteolysis | yes | yes | 4 | 5 |
| 15 | Other: intracellular signal transduction | no | no | 3 | 13 |
| 16 | Other: syntaxin binding | no | no | 4 | 7 |
| 17 | Steroid / progesterone metabolism | yes | yes | 3 | 2 |
| 18 | Other: protein tyrosine kinase activity | no | no | 6 | 6 |
